# Tumor microenvironment acidosis favors pancreatic cancer stem cell properties and *in vivo* metastasis

**DOI:** 10.1101/2024.06.14.599032

**Authors:** Michala G. Rolver, Juan C. Roda, Yifan Dai, Mette Flinck, Renata Ialchina, Julie Hindkær, Rigmor T. Dyhr, August N. Bodilsen, Nanditha S. Prasad, Jonathan Baldan, Jiayi Yao, Albin Sandelin, Luis Arnes, Stine F. Pedersen

**Affiliations:** Section for Cell Biology and Physiology, Department of Biology, Faculty of Science, University of Copenhagen, Denmark; Biotech Research and Innovation Centre, University of Copenhagen, Denmark; The Bioinformatics Centre, Department of Biology, University of Copenhagen, Denmark

**Keywords:** pancreatic ductal adenocarcinoma, PDAC, pancreatosphere, pH regulation, CSC, orthotopic

## Abstract

The acidic tumor microenvironment favors cancer aggressiveness via incompletely understood pathways. Here, we asked whether acidic environments select for cancer stem cell (CSC) properties. Bulk RNA-seq of Panc-1 human pancreatic cancer cells adapted to extracellular pH 6.5 revealed upregulation of CSC markers including CD44, EpCam, Nestin and aldehyde dehydrogenases, and CSC pathway enrichment. We therefore assessed CSC characteristics of acid-adapted (AA) and non-adapted (Ctrl) PaTu8988s and MiaPaca-2 pancreatic cancer cells. Compared to Ctrl, AA cells exhibited increased ALDH- and β-catenin activity and pancreatosphere-forming efficiency, classical CSC characteristics. Panc-1, PaTu8988s and MiaPaCa-2 AA cells differed in CSC marker expression, and AA cells did not exhibit typical flow cytometric CSC populations. However, single-nucleus sequencing identified the acid adaptation-induced emergence of a population with clear CSC characteristics. Finally, in an orthotopic mouse model, AA Panc-1 cells drove strongly increased aggressiveness and liver metastasis compared to Ctrl cells.

We conclude that acid-adaptation of pancreatic cancer cells leads to enrichment of a CSC phenotype with unusual traits, providing new insight into how acidic tumor microenvironments favor cancer aggressiveness.

## Introduction

Pancreatic cancer is one of the most lethal cancers globally, with a 5-year survival rate of less than 10% (Neoptolemos *et al*, 2018). Despite important recent developments (Rojas *et al*, 2023), first-line treatment for pancreatic cancer remains chemotherapy and radiation, yet these only increase survival by weeks or a few months (Neoptolemos *et al*, 2018). The global burden of pancreatic cancer has greatly increased in the past decades and this is likely to continue as the population ages (GBD 2017 Pancreatic Cancer Collaborators, 2019), thus highlighting the need for improved understanding of the molecular mechanisms of this disease and translation of this to new therapeutic strategies. During their growth, some regions in solid tumors develop extensive acidosis, resulting from the combination of elevated metabolic acid production and insufficient vascularization (Swietach *et al*, 2023). This is also the case for pancreatic cancers (Kimbrough *et al*, 2015), in which this has been suggested to be particularly important, given the extreme acid-base transport physiology of the pancreatic duct (Pedersen *et al*, 2017). Together with other tumor microenvironment (TME) traits such as hypoxia, acidosis plays a key role in cancer progression (Swietach *et al*, 2023)(Boedtkjer & Pedersen, 2020; Corbet & Feron, 2017). We and others have shown that cancer cells adapted to growth at an extracellular pH (pH_e_) corresponding to that measured in acidic tumor regions, develop aggressive traits such as increased 3D growth and adhesion-independent colony formation (Czaplinska *et al*, 2023) and increased invasiveness and metastatic potential (Czaplinska *et al*, 2023)(Corbet *et al*, 2020)(Moellering *et al*, 2008). Notably, acidic growth also rewired cancer cell metabolism toward increased reliance on oxidative phosphorylation (OXPHOS) and fatty acid β-oxidation (Michl *et al*, 2022; Rolver *et al*, 2023)(Corbet *et al*, 2014)(Lamonte *et al*, 2013), and profoundly altered their iron- and redox homeostasis (Dierge *et al*, 2021).

Cancer stem cells (CSCs), also referred to as Tumor-initiating cells (TICs), were first identified in hematological cancers and since discovered in essentially all cancers. While their existence is no longer controversial, their properties, origin, and essential nature remain incompletely understood (Batlle & Clevers, 2017)(Vermeulen *et al*, 2008). It is increasingly appreciated that CSCs are highly plastic and can derive from de-differentiation of non-stem cells such as progenitor cells or more differentiated tumor cells (Hermann & Sainz, 2018)(Batlle & Clevers, 2017)(Vermeulen *et al*, 2008). These processes and the maintenance of the CSC state, are driven by incompletely understood signals that, at least in part, derive from the CSC niche – a distinct part of the TME (Plaks *et al*, 2015). In pancreatic cancer, CSCs have been assigned a key role in the initiation, aggressiveness, and chemotherapy resistance of the disease (Hermann & Sainz, 2018; Patil *et al*, 2021)(Li *et al*, 2007)(Hermann *et al*, 2007). Interestingly, CSCs are metabolically distinct from other cancer cells, with a shift toward mitochondrial metabolism (Vlashi *et al*, 2011; Sancho *et al*, 2015)(Raggi *et al*, 2021).

Specifically pancreatic CSCs were shown to exhibit upregulated OXPHOS and to strongly rely on mitochondrial metabolism (Courtois *et al*, 2021)(Sancho *et al*, 2015), and JAK/STAT3-driven increased fatty acid β-oxidation was recently shown to be essential for CSC phenotype and chemoresistance in breast cancer cells (Wang *et al*, 2018). This metabolic phenotype resembles that which we recently demonstrated in acid-adapted pancreatic cancer cells (Rolver *et al*, 2023). Furthermore, similar to CSCs, acid-adapted cancer cells exhibited increased adhesion-independent colony formation (Czaplinska *et al*, 2023).

In this work, we tested the hypothesis that acid adaptation drives CSC properties. We show, using three different pancreatic cancer cell lines, that adaptation to growth in an acidic microenvironment is associated with upregulation of CSC properties such as increased ALDH- and β-catenin activity and enhanced pancreatosphere formation capacity, and emergence of a stem cell population as seen by single cell RNA sequencing, but with an unusual marker expression pattern compared to previously described pancreatic CSCs. Furthermore, acid-adapted Panc-1 cells drove aggressive metastatic growth *in vivo*. We suggest that acid adaptation-induced upregulation of CSC phenotypic traits contributes to the aggressive cancer growth favored by the acidic tumor microenvironment.

## Results

### CSC genes, ALDH- and β-catenin activity are upregulated in acid-adapted pancreatic cancer cells

In earlier work, we studied the impact of acidic tumor niches on the phenotype of cancer cells from different tissue origins, by adapting them to growth in acidic conditions (Fig. 1A; (Yao *et al*, 2020; Rolver *et al*, 2023). In light of the growing awareness of the role of CSCs in pancreatic cancer (Hermann & Sainz, 2018; Patil *et al*, 2021), we revisited our RNA-seq analysis of Panc- 1 pancreatic cancer cells to ask if CSC markers were upregulated in acid adapted (AA, pH 6.5) cells compared to cells grown at extracellular pH (pH_e_) 7.4: we extracted log_2_ fold change (FC) values for the significantly differentially expressed genes in Panc-1 cells and ranked them from the most upregulated to most downregulated compared to Ctrl cells (Fig. 1B). On this ranked set, we mapped the position of reported pancreatic CSC markers (Fig. 1B). These include cluster of differentiation (CD) *CD44, CD24*, and epithelial-specific antigen (*ESA*, aka EpCAM) (Li *et al*, 2007), *CD133*, C-X-C chemokine receptor type 4 (*CXCR4*) (Hermann *et al*, 2007), SRY-box transcription factor-9 (*SOX9*), *NANOG*, the ABC efflux transporter *ABCG2* (Nallasamy *et al*, 2021), the intermediate filament protein Nestin (*NES*), c-Met (Ishiwata *et al*, 2018), the tetraspanin *CD9*, the glutamine transporter ASCT2 (*SLC1A5*), the lactate-H^+^ cotransporter MCT1 (*SLC16A1*) (Wang *et al*, 2019)(Jang *et al*, 2022), and the long non-coding RNA HOX antisense intergenic RNA (*HOTAIR*) (Wang *et al*, 2017). We also mapped the aldehyde dehydrogenase (ALDH) genes ALDH1A3, *ALDH3B1*, *ALDH5A1* and *ALDH6A1,* given the key roles of ALDH activity for CSCs, including PDAC CSCs (Kim *et al*, 2011). This analysis revealed a strong tendency for CSC genes to be upregulated in AA Panc-1 cells (Fig. 1B). On the other hand, the Yamanaka stem cell factors, i.e. Krüppel-like factor 4 (*KLF4*), octamer-binding transcription factor (*OCT)3/4, SOX2,* and *MYC* (Takahashi & Yamanaka, 2006), tended to be downregulated in AA cells (Fig. 1B). The association of acid adaptation with increased stemness was further supported by gene set enrichment analysis (GSEA) of all significantly upregulated genes in Panc-1-AA cells relative to Ctrl, using the SigDb database of gene sets. The acid-adapted Panc-1 gene set was clearly enriched for genes upregulated in glioma- and liver cancer stem cells, whereas this overlap was not observed for a set of genes upregulated in leukemia stem cells (Fig. 1C), and was also not seen for non-cancer embryonic stem cells (ESCs, Suppl. Fig. 1A).

**Figure 1.**
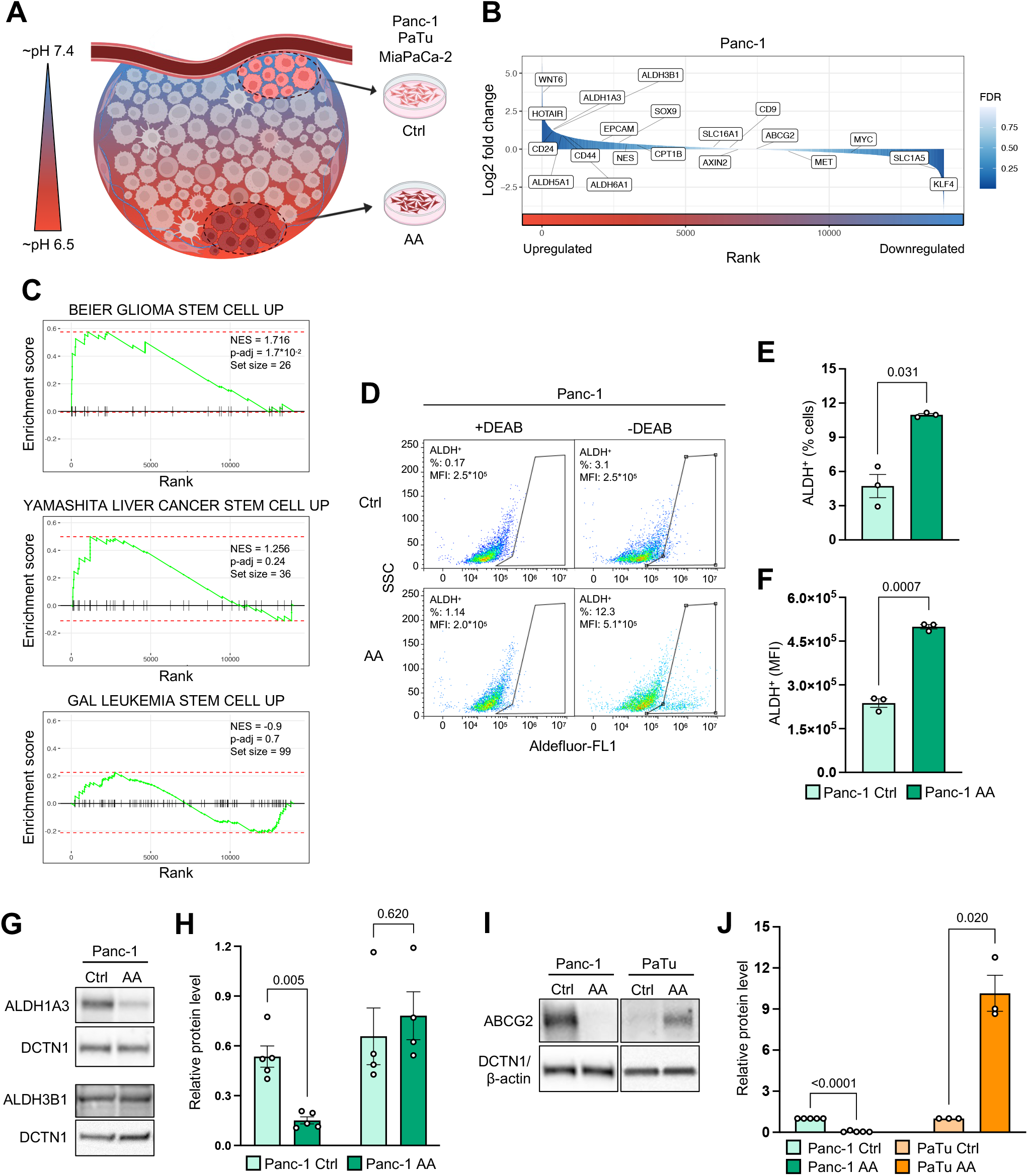
Acid-adapted pancreatic cancer cells exhibit increased mRNA expression of stemness-associated genes. (**A**) Conceptual illustration of the experimental setup. Panc-1, PaTu-8988 (PaTu) or MiaPaCa- 2 cells were grown in growth medium mimicking tumor areas with either physiological or acidic pH_e_ (pH_e_:∼7.4, Ctrl or ∼6.5, acid-adapted, AA). Ctrl and AA cells were cultured in parallel for 1 month to allow AA cells to adapt to the acidic pH_e_. (**B**) Fold change-based ranking of genes expressed in AA Panc-1 cells. The x-axis is the rank assigned to each gene based on gene expression fold change (AA vs. Ctrl) shown as y-axis in log2 scale. The color intensity corresponds to differential expression significance, expressed as false discovery rate (FDR) in -log10 scale. Genes associated with stemness are labeled in the plot. (**C**) Gene set enrichment analyses (GSEA) based on acid adaptation-ranked gene list and selected stemness-relevant gene sets in SigDB database. The X axis shows genes ranked by their log2FC (AA vs Ctrl). The Y-axis shows the enrichment score, reflecting the over-representation level of the chosen gene set in the up- or downregulation parts of the ranked acid adaptation gene list (X axis). Vertical lines across each panel indicate the occurrence of genes from the selected SigDB set in the ranked acid adaptation gene list. The overall normalized enrichment score (NES), associated *FDR* value (p-adj) and set size are given in each panel. (**D**) Representative flow cytometry plots of aldehyde activity in Panc-1 Ctrl and AA cells measured employing the ALDEFLUOR^TM^ assay in presence or absence of diethylaminobenzaldehyde (DEAB). The numbers inside the plots denote the ALDH^+^ cells in percent (%) and mean fluorescent intensity (MFI). (**E-F**) Quantification of ALDH activity expressed as percent ALDEFLUOR^TM^ positive cells (ALDH^+^) (**E**) and MFI of the ALDH^+^ population (**F**). (**G-H**) Representative Western blots (**G**) and corresponding quantification of protein levels (**H**) of ALDH1A3 and ALDH3B1 in Panc-1 Ctrl and AA cells. Dynactin1 (DCTN1) was used as loading control. (**I-J**) Representative Western blots (**I**) and corresponding quantification of protein levels (**J**) of ABCG2 in Panc-1 and PaTu Ctrl and AA cells. DCTN1 and β-actin were used as loading control for Panc-1 and PaTu, respectively. For panels (**E**, **F**, **H**, **J**) a paired two-tailed student’s t-test was performed for each individual cell line, AA vs Ctrl.

In agreement with the upregulation of multiple ALDH isoforms in AA Panc-1 cells (Fig. 1B), both the fraction of cells with ALDH activity and the mean fluorescence intensity (MFI) of the ALDH positive population (ALDH+) were upregulated in AA Panc-1 cells compared to Ctrl cells (Fig. 1D-F), a strong indication of increased CSC characteristics (Kim *et al*, 2011)(Xu *et al*, 2015). Surprisingly, neither ALDH1A3 nor ALDH3B1 were upregulated at the protein level in AA Panc-1 cells compared to Ctrl cells (Fig. 1G-H), suggesting that another upregulated ALDH isoform (Fig. 1B), or post-translational regulation of ALDH activity, is responsible for the marked upregulation of ALDH activity in AA cells.

To further study the possible link between acid adaptation and pancreatic cancer stemness, we adapted two additional human pancreatic cancer cell lines, PaTu8988s (PaTu) and MiaPaca-2, to grow at pH 6.5 (AA) and pH 7.6 (Ctrl) for 1 month (as in Fig. 1A). mRNA levels of *ALDH1A3* were unaffected by acid adaptation in PaTu cells but strongly upregulated by this treatment in MiaPaca-2 cells (Suppl. Fig. 1B-C).

Another classical CSC assay is the side population (SP) assay, which is based on the upregulation in CSCs of Hoechst extrusion via ATP-binding cassette (ABC) transporters, primarily but not exclusively, ABCG2 (Bcrp1) (Zhou *et al*, 2001). The ABCG2 protein level was increased in AA PaTu and AA MiaPaCa-2 cells yet downregulated in AA Panc-1 cells (Fig. 1I-J, Suppl. Fig. 1D-E), arguing against a general ABCG2 upregulation in AA cells.

These results show that acid adapted pancreatic cancer cell lines, to a variable extent, exhibit gene expression changes and phenotypic traits characteristic of CSCs.

### Acid-adapted pancreatic cancer cells exhibit increased β-catenin signaling

Increased signaling via the Wnt-β-catenin pathway (Fig. 2A) contributes to the phenotypic traits of many CSCs, including pancreatic CSCs (Wu *et al*, 2019). We noted that *WNT6* was one of the most highly upregulated genes in AA Panc-1 cells in our RNA-seq analysis (Fig. 1B), and KEGG analysis of WNT signaling showed differential regulation of multiple pathway components (Suppl. Fig. 2A). Western blotting for Wnt6 showed that WNT6 protein levels were increased in Panc-1 AA cells compared to Ctrl cells (Fig. 2B-C). The β-catenin protein level was increased in AA cells compared to Ctrl, consistent with increased Wnt signaling in the AA cells (Fig 2B-C). Inhibitory GSK-3β phosphorylation at Ser9 was not substantially altered, which was expected if GSK-3β inhibition occurs via the Wnt pathway, in which GSK-3β is instead inhibited by sequestering (Metcalfe & Bienz, 2011) (Fig. 2B-C). Consistent with the increased β-catenin protein level, also β-catenin transcriptional activity, as assessed using the TOPFlash luciferase reporter, was increased in AA Panc-1 cells compared to Ctrl (Fig. 2D). Wnt signaling can also activate c-Jun N-terminal kinase (JNK) signaling (Lien & Fuchs, 2014). Activating phosphorylation of p46JNK was indeed increased in AA cells compared to Ctrl cells (Fig. 2E-F). Interestingly, also total p46JNK levels were increased, and consequently, the p-p46JNK/total p46JNK ratio was not significantly increased (Fig. 2E-F). Also the total p54JNK level was increased in AA cells, but p-p54JNK was not altered (Suppl. Fig. 2B-C).

**Figure 2.**
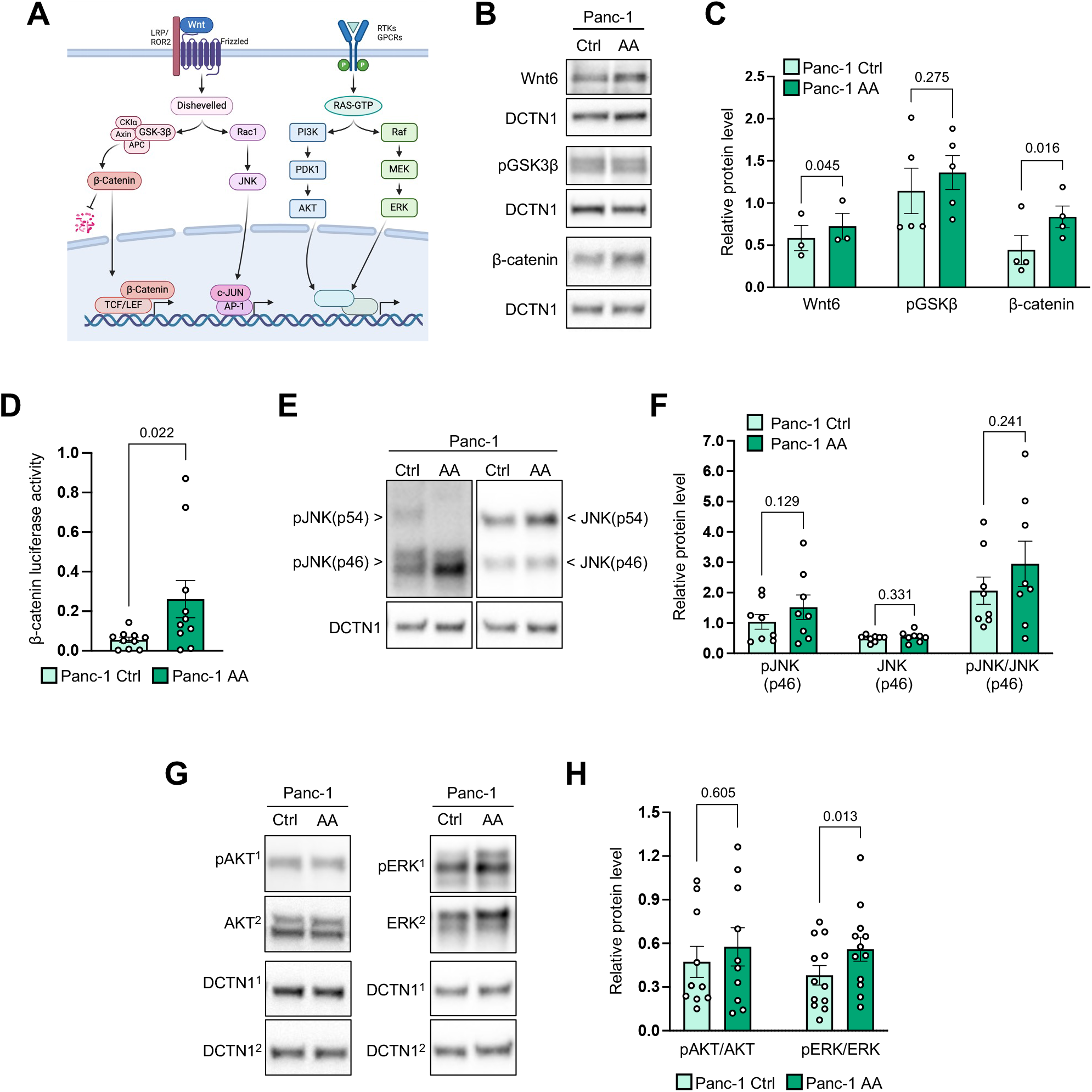
Acid-adapted pancreatic cancer cells exhibitincreased β-catenin signaling. (**A**) Simplified illustration of the canonical and non-canonical Wnt pathways (first and second pathways from the left, respectively) and receptor-mediated AKT and ERK signaling pathways (third and fourth pathways from the left, respectively). Wnt signaling can regulate stem cell pluripotency and cell fate (canonical) and cytoskeleton and cell migration (non-canonical); AKT and ERK signaling can regulate proliferation and survival. (**B-C**) Representative Western blots (**B**) and corresponding quantification of protein levels (**C**) of Wnt6, pGSK3β and β-catenin in Panc-1 Ctrl and AA cells. DCTN1 was used as loading control. (**D**) Relative β-catenin activity-mediated Firefly luciferase signal normalized to pRL-TK-mediated Renilla luciferase signal in Panc-1 Ctrl and AA cells measured by dual luciferase assay. Data was log-transformed prior to testing for significance to improve normal distribution. (**E-H**) Representative Western blots (**E**, **G**) and corresponding quantification of protein levels of p-p46JNK/p46JNK (**F**) and pAKT/AKT and pERK/ERK (**H**) in Panc-1 Ctrl and AA cells. DCTN1 was used as loading control. In (**G**), the 1 and 2 in superscript denote the pairing between protein of interest and its corresponding loading control. For panels (**C**, **D**, **F**, **H**) a paired two- tailed student’s t-test was performed, AA vs Ctrl.

AKT- and extracellular signal regulated kinase (ERK1/2) signaling is hyperactivated in pancreatic cancers downstream of activating KRAS mutations (Eser *et al*, 2013). Interestingly, while AKT activity was unaltered, ERK activity, assessed as its activating phosphorylation, was increased in AA Panc-1 cells compared to Ctrl cells (Fig. 2G-H).

These results show that Wnt signaling to β-catenin and JNK, as well as ERK1/2 activity, are upregulated in acid-adapted Panc-1 cells.

### Acid-adapted pancreatic cancer cells exhibit increased pancreatosphere-forming capacity

Encouraged by these findings, we next studied the ability of acid-adapted pancreatic cancer cells to form pancreatospheres – a classical measure of stemness (Wang *et al*, 2013). Cells were thinly seeded in suspension in pancreatosphere medium, grown for 7 days, and pancreatospheres (first generation) were imaged, analyzed by counting or image analysis of pancreatosphere area. Pancreatospheres were then dissociated, and the process repeated for up to three generations (Fig. 3A). Both Ctrl and AA PaTu cells were capable of growing lumen-forming, organoid-like colonies under these conditions (Fig. 3B). Strikingly however, these structures were significantly larger in AA cells than in Ctrl cells (Fig. 3B) and the total number of colonies (pancreatosphere forming efficiency, PFE) was significantly increased for AA cells compared to Ctrl cells, for both the first and second generation of pancreatospheres seeded (Fig. 3C).

**Figure 3.**
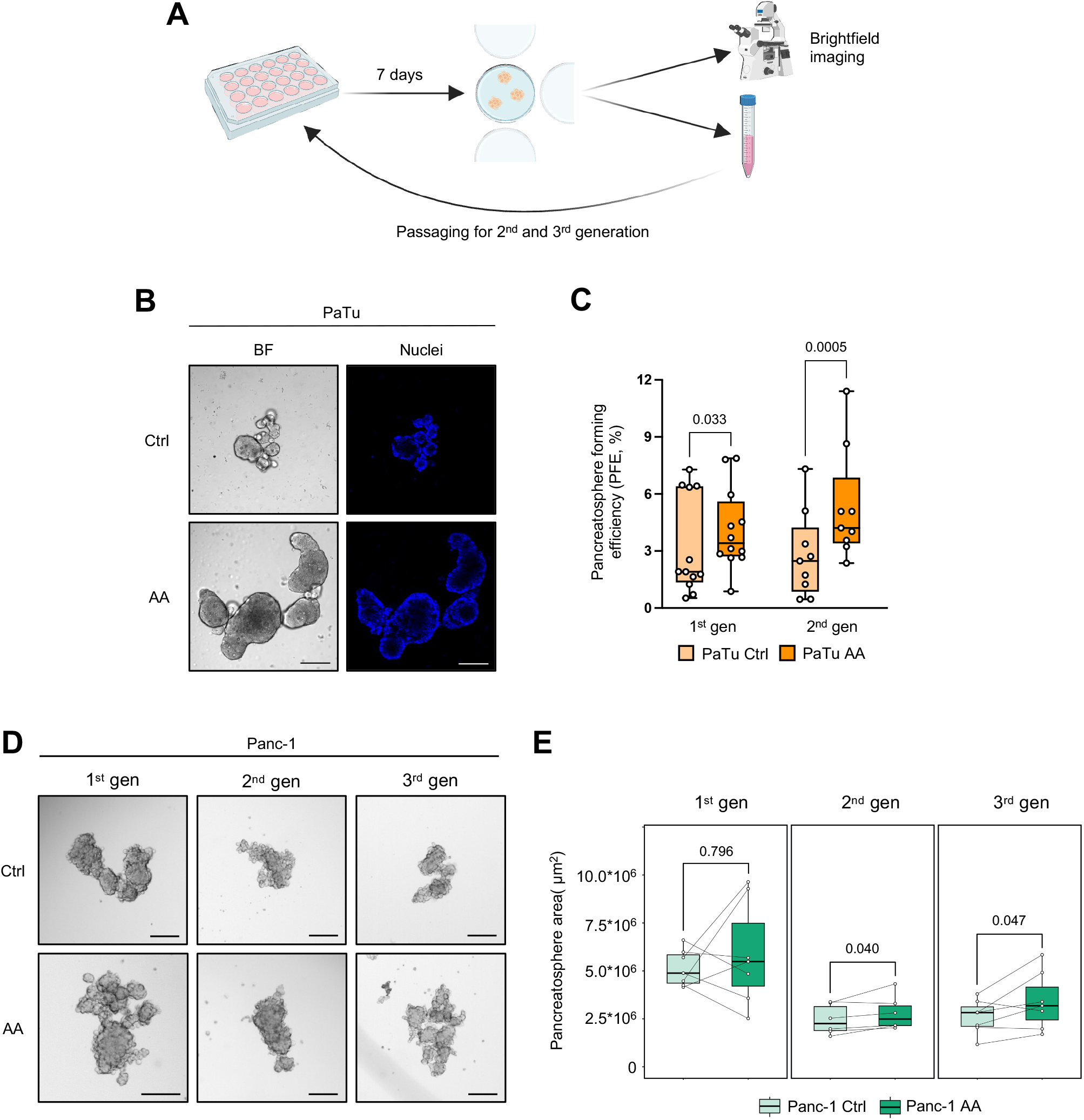
Acid-adapted pancreatic cancer cells exhibit increased pancreatosphere-forming capacity. (**A**) Illustration of the workflow of the pancreatosphere growth assay. (**B**) Representative brightfield (BF, left) and fluorescence (right) microscopy images of 1^st^ generation PaTu Ctrl and AA pancreatospheres. Scale bar: 200 µm. (**C**) Box plot of the pancreatosphere forming efficiency in percent for 1^st^ and 2^nd^ generation (gen) of PaTu Ctrl and AA pancreatospheres. In each biological replicate (points on plot), 2-3 individual wells were analyzed. Horizontal line denotes median, and upper and lower lines indicate max. and min. values, respectively. (**D**) Representative BF microscopy images of Panc-1 Ctrl and AA pancreatospheres from 1^st^, 2^nd^ and 3^rd^ generation (gen). Scale bar: 200 µm. (**E**) Box plot of the mean pancreatosphere area (µm^2^) for each generation of Panc-1 Ctrl and AA pancreatospheres. In each biological replicate (points on plot), 3-7 individual wells were analyzed. The lines connecting the points within a generation denote the pairs of Ctrl and AA pancreatospheres in each biological replicate. For (**C**, **E**) a paired two-tailed student’s t-test was performed for AA vs Ctrl within each generation.

A similar pattern was seen for Panc-1 cells, which were re-seeded for a total of 3 generations and pancreatosphere area was determined by semi-automated image analysis (Fig. 3D-E). For these cells, the first generation of pancreatospheres did not differ between Ctrl and AA cells. However, in both the second and third generation, AA pancreatospheres occupied a significantly larger area than those formed by Ctrl cells, showing that the capacity for pancreatosphere formation is maintained over several generations (Fig. 3D-E). Finally, also MiaPaCa-2 cells, followed for 2 generations, showed increased pancreatosphere-forming capacity when acid-adapted (Suppl. Fig. 3A-B).

Collectively, these results show that acid-adapted pancreatic cancer cells exhibit increased self-renewal capacity, an important feature of CSCs.

### CSC markers are upregulated in subpopulations of acid-adapted pancreatic cancer cells

A wide variety of CSC populations and CSC surface markers have been reported in pancreatic cancer (Ishiwata *et al*, 2018)(Jaiswal *et al*, 2012; Kallifatidis *et al*, 2009)(Nimmakayala *et al*, 2021)(Li *et al*, 2007)(Wang *et al*, 2019). Furthermore, the possible contributions of individual CSC subpopulations to disease aggressiveness are essentially unknown (Hermann & Sainz, 2018; Patil *et al*, 2021)(Zhao *et al*, 2023). To gain insight into which stem cell markers are expressed by acid-adapted pancreatic cancer cells, we studied the expression of the Yamanaka stem cell factors, KLF4, SOX2, and Oct4, and the most widely reported pancreatic cancer CSC markers (Fig. 4A), by Western blotting, qPCR, and flow cytometry. Consistent with our RNA-seq data (Fig. 1B), the Yamanaka factors *OCT4* and *SOX2* mRNA levels were downregulated in AA Panc-1 cells (Suppl. Fig. 4A). At the protein level, SOX2 was increased in AA Panc-1 but decreased in AA PaTu cells, whereas KLF4 was decreased in AA Panc-1 cells, (marginally) increased in AA PaTu cells, and significantly increased in MiaPaCa-2 cells compared to Ctrl (Suppl. Fig 4B-C).

**Figure 4.**
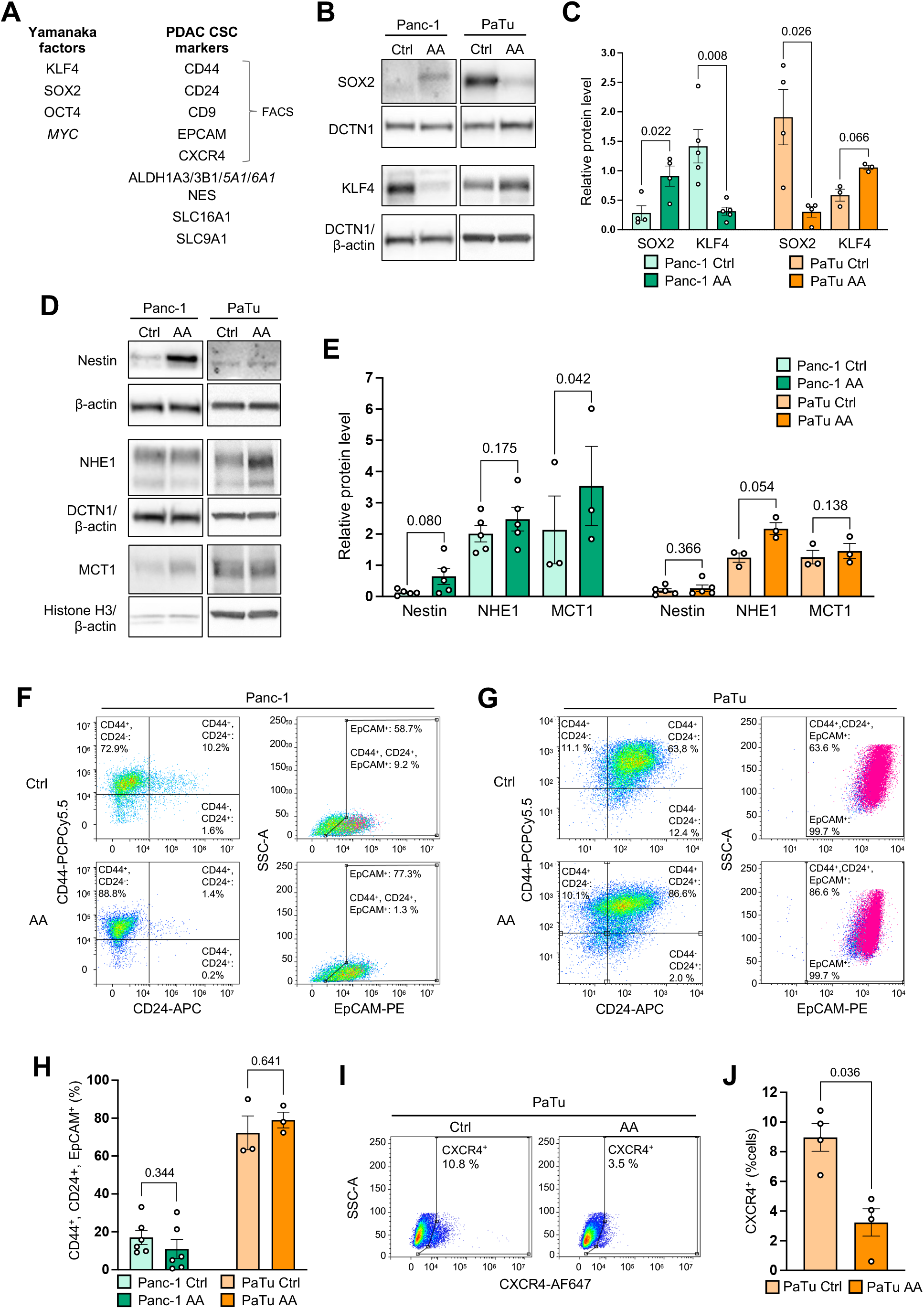
CSC markers are upregulated in subpopulations of acid-adapted pancreatic cancer cells. (**A**) Table of selected stem cell markers divided into the classical Yamanaka factors (left side) and PDAC-specific CSC factors (right side). Genes in italics have not been experimentally investigated. (**B**-**E**) Representative Western blots (**B, D**) and corresponding quantification of protein levels (**C, E**) of SOX2 and KLF4 (**B**-**C**) and Nestin, NHE1 and MCT1 (**D-E**) in Panc-1 and PaTu Ctrl and AA cells. DCTN1, β-actin or histone-H3 were used as loading controls as indicated. Where two loading controls are noted, the upper relates to Panc-1 cells and the lower to PaTu cells. (**F**-**H**) Representative flow cytometry scatter plots (**F, G**) and quantification (**H**) of cell surface expression of CD44, CD24 (left side), and EpCAM (right side) in Panc-1 (**F, H**) and PaTu (**G, H**) Ctrl and AA cells as indicated. Triple positive (CD44^+^, CD24^+^, EpCAM^+^) cells are marked in pink in the EpCAM plot. (**I-J**) Representative flow cytometry scatter plots (**I**) and quantification (**J**) of cell surface expression of CXCR4 in PaTu Ctrl and AA cells. For panels (**F**, **G**, **I**), the numbers inside the plots denote the percent positive cells of the specified marker(s). In panels (**C**, **E**, **H**, **J**), a paired two-tailed student’s t-test was performed for each individual cell line, AA vs Ctrl.

We next studied the pancreatic CSC markers Nestin and SLC16A1 (MCT1). We also included the net acid extruding transporter Na^+^/H^+^ exchanger-1 (NHE1, SLC9A1), a key regulator of cellular pH during acidosis (Pedersen & Counillon, 2019), changes in the activity of which have been associated with regulating transitions between stemness and differentiation (Liu *et al*, 2023). The protein levels of these three genes were either unaltered or modestly increased in acid adapted cells compared to their respective controls in all three cell lines (Fig. 4D-E, Suppl. Fig. 4D-I). Finally, we assessed mRNA levels of other genes reported to be differentially expressed in pancreatic CSCs including *KLF9, ING4, PSMD11,* and *HOTAIR*. With the exception of PSMD11, which was increased in PaTu cells, none of these mRNAs showed altered expression in PaTu and MiaPaCa-2 cells (Suppl. Fig. 4J).

Next, we explored potential CSC subpopulations using flow cytometry of CSC surface markers in Ctrl and AA Panc-1 and PaTu cells. The most widely studied pancreatic CSC population is reported to be CD44^+^/CD24^+^/EpCaM^+^. This population comprised about 15% of Panc-1 cells and a very high fraction – around 75% – of PaTu cells, and did not change upon acid adaptation in any of the two cell lines (Fig. 4F-H). Finally, surface levels of CD9, another described CSC marker, was not changed by acid adaptation of Panc-1 and PaTu cells (Suppl. Fig. 4K-L).

These results show that protein levels of some classical stem cell and CSC markers and net acid-extruding transporters linked to stemness are upregulated by acid-adaptation. However, others are downregulated, and we do not observe a general pattern of protein level upregulation of CSC markers by acid adaption.

### Acid adaptation leads to distinct cellular subpopulations with upregulated stem cell markers

Based on the results above, we hypothesized that the fraction of stem-cell-like cells would increase after acid adaptation but that its identification might require analysis at single cell level, because cells may employ different strategies to adapt to an acid microenvironment. To investigate this, isogenic human Panc-1 cells were randomly split into control and acid adaptation groups with three replicates in each group. AA samples were cultured under acidic conditions (pH 6.7) for 4 weeks, while Ctrl samples were maintained at constant physiological pH of 7.6 for the same time. At the end of the adaptation, nuclei from each cell group were extracted and subjected to single cell RNA sequencing using the 10x Genomics snRNA-seq workflow. One sample in each group was harvested twice, resulting in two technical AA and Ctrl replicates. Sequenced reads were mapped to the human transcriptome, followed by quality control and analysis using Seurat (Hao *et al*, 2024) (see Materials and Methods). After quality filtering, we recovered 2143 to 5930 cells per replicate (Fig. S5a)

UMAP visualization of all cells showed three major cell clusters, where one primarily was composed of Ctrl cells and two were composed of AA cells. We denoted these the Ctrl, AA1 and AA2 clusters, respectively (Fig 5A-B). While the Ctrl cluster was evenly populated by cells from Ctrl replicates (including the Ctrl technical replicate), only the part of the AA1 cluster closest to the Ctrl cluster was populated by cells from all AA replicates. The rest of the AA1 cluster could be divided into two parts, of which the first was populated by cells from AA replicate 1 and 2, and the other by cells from AA replicate 3 (and its technical replicate). Finally, the AA2 cluster was dominated by the AA replicate 1. A high proportion of G0/G1 phase cells is a hallmark of some CSCs (Davis *et al*, 2019). We found that the average proportion of G0/G1 cells increases by 4.2-fold in AA1 and 6.7-fold in AA2 cells compared to Ctrl (Fig. S5B). Interestingly, the highest proportion of G0/G1 cells occurred in, or close to, the part of the AA1 cluster that was populated by all AA replicates (Fig. 5A, Fig. S5C)

**Figure 5.**
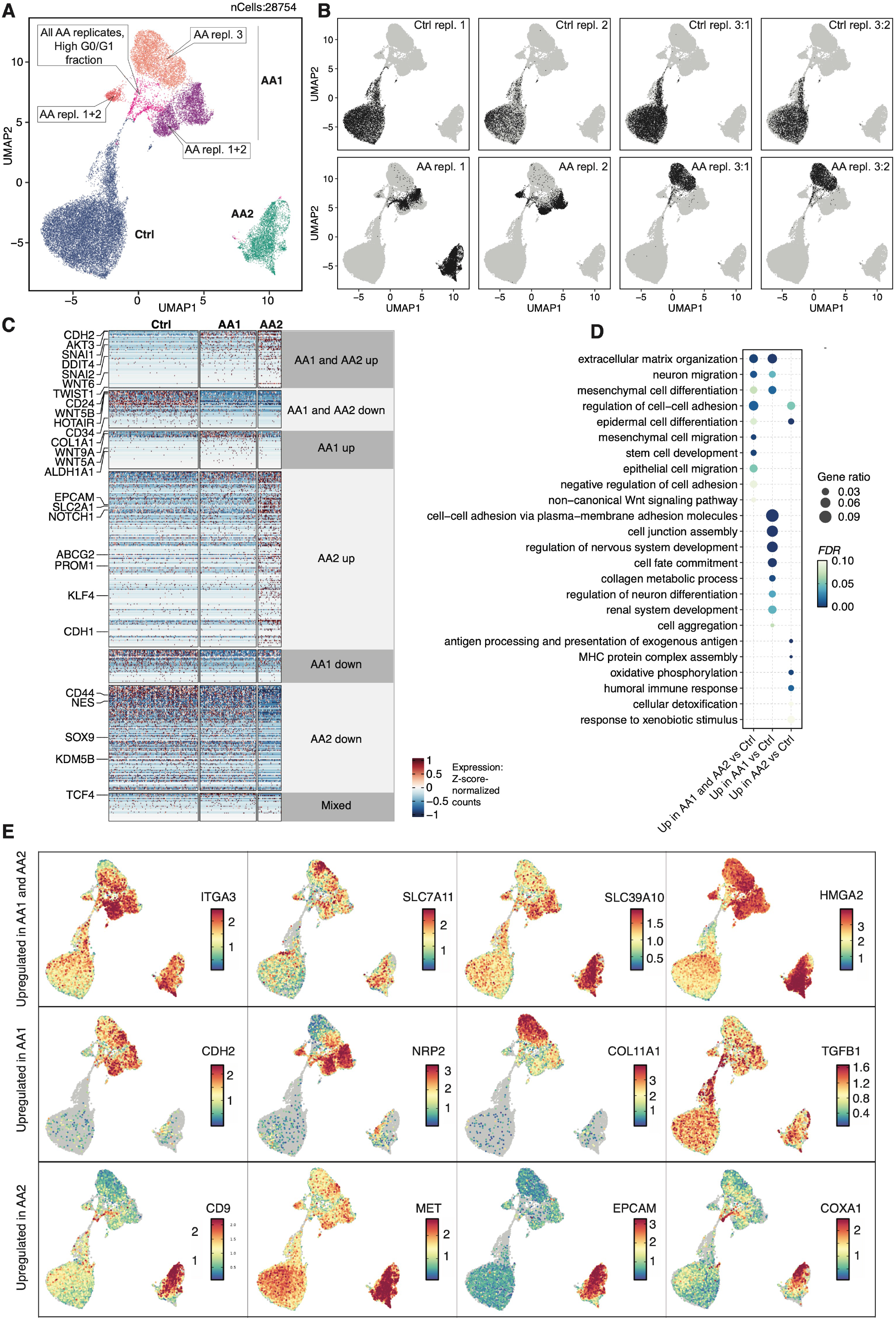
Single nuclei RNA-seq indicates the existence of several acid-adapted expression phenotypes, each expressing stem cell marker genes. **(A)** UMAP visualization of all RNA-seq replicates from single nuclei. The x- and y-axis correspond to UMAP1 and 2, respectively. Each dot visualizes RNA-seq data in one nucleus. Three replicates of AA cells (AA 1-3) and cells grown at static pH 7.6 (Ctrl 1-3) were sequenced; one replicate in each group was sequenced twice (technical replicates: AA3:1-2 and Ctrl3:1-2). Colors show UMAP clusters: the AA1 cluster was divided into colored subclusters with callouts summarizing subcluster features (also shown in panel **B**). **(B)** UMAP visualization of individual RNA-seq replicates. Each subpanel shows the UMAP from panel **A** where cells from a given replicate are colored black and remaining cells are colored gray. Replicates 3:1 and 3:2 denote the technical replicates mentioned in (**A**) within each group. **(C)** Heat map visualization of differentially expressed genes between AA1, AA2 and Ctrl clusters. Each row corresponds to one gene, and each dot corresponds to one cell, colored by Z-score normalized expression. Bars on top show clusters from the UMAP in (**A**). Rows are divided into groups with similar expression patterns (e.g. upregulation in AA1 vs Ctrl). Names of stemness-associated genes are shown to the left. (**D**) Over-representation of selected GO terms in DEGs upregulated in the AA1 and/or AA2 cluster vs Ctrl. Rows show GO terms. Columns show DEG groups. Dot size indicates gene ratio, defined as the proportion of GO term genes in each gene group while color indicates over-representation significance (*FDR*). **(E)** UMAP visualization of expression of key genes upregulated in AA1, AA2 or both vs Ctrl., as discussed in the main text. UMAPs as in (**A**) but colored by the expression of the indicated gene. UMAPs are organized by groups of genes upregulated in both, or only one, AA cluster vs. ctrl. Also see Suppl. Fig. 5D.

To further explore whether stem cell markers and associated genes were enriched in AA1, AA2 or both vs Ctrl, we performed differential expression (DE) analysis. Fig 5C shows a heatmap of DE genes (log_2_FC > 1, *FDR* < 0.05). We noted that several of the upregulated genes in these clusters are associated with stemness (gene names highlighted in Fig 5C). Agreeing with this, gene ontology (GO) analysis of genes upregulated in one or both clusters vs Ctrl also showed overrepresentation of functional categories related to stemness and CSCs: for example GO terms for ECM, neural migration, mesenchymal cell differentiation, regulation of cell adhesion and non-canonical Wnt pathway GO terms were enriched in upregulated genes in both clusters. Additionally, AA1-only upregulated genes were enriched for cell fate commitment, cell junction assembly, and AA2-only upregulated genes were enriched for several immune- and xenobiotic response terms, and for oxidative phosphorylation (Fig. 5D shows selected terms, Suppl. table 1 shows all terms analyzed). We discuss a handful of DE genes associated with these terms below. Expression data for these genes is shown in Fig 5E and Fig. S5D

Key upregulated genes in both AA1 and AA2 included *SEMA3A*, associated to glioblastoma stem cell proliferation and invasion (Higgins *et al*, 2020), *ITGA3*, encoding integrin-α3 and shown to drive aggressive disease in pancreatic cancer (Liu *et al*, 2021)(Jiao *et al*, 2019), *SLC39A10*, encoding the zinc transporter ZIP10, which is associated with aggressive cancers, migration, inflammation, and anti-apoptosis and is a direct Myc target (Ren *et al*, 2023; Ma *et al*, 2021; Miyai *et al*, 2014) and also identified through our previous bulk RNA-seq analysis of acid-adapted cells (Rolver *et al*, 2023), *SLC7A11*, encoding the cystine-glutamate transporter xCT, which is overexpressed in many cancers, and protects cancer cells and stem cells from oxidative stress and ferroptosis (Yan *et al*, 2023)(Wu *et al*, 2022) and *HMGA2*, a regulator of epithelial-mesenchymal transition (EMT) and an important stemness driver (Wu *et al*, 2022; Li *et al*, 2017).

AA1-specifically upregulated genes included *CDH2*, encoding N-cadherin, a widely known EMT marker, *NRP2*, a semaphorin receptor implicated in EMT and metastasis (Xu *et al*, 2023) (notably, its ligand *SEMA3A*, associated with tumor growth, was also upregulated in the AA1 cluster), and a large number of genes associated to ECM organization e.g. *COL11A1, COL5A2, COL1A1 and TGFB1*. Other stem cell- or EMT-associated genes in the AA1 cluster included *VEGFC*, *ERBB4*, as well as *GLI2*, a major effector in Hedgehog signaling, which is a well-established driver of pancreatic cancer stemness (Lee *et al*, 2008).

Remarkably, genes upregulated specifically in the AA2 cluster included the established pancreatic cancer stem cell markers, *CD9* (Wang *et al*, 2019) which was also found in our bulk RNA-seq analysis (Fig. 1b), *MET*, encoding the receptor tyrosine kinase c-MET (Ishiwata *et al*, 2018; Pothula *et al*, 2020), and *EPCAM* (Gires *et al*, 2020; Ishiwata *et al*, 2018). This strongly indicates that our inability to detect these by other methods reflected that only a fraction of the acid-adapted cells showed upregulation of these CSC markers. Furthermore, a large number of genes associated with oxidative phosphorylation and/or mitochondria were upregulated in AA2, e.g. *COX6A1, COX6C, COX7C, COX4I1*. This is consistent with the important role of oxidative phosphorylation in CSCs (Sancho *et al*, 2015)(Courtois *et al*, 2021), and with data from us and others pointing to the dependence of acid-adapted cancer cells on oxidative phosphorylation (Rolver *et al*, 2023; Michl *et al*, 2022).

Overall, these results indicate that acid adaptation may lead to different transcriptional phenotypes for subsets of cells, some of which always or often occur (eg the shared region of AA1) and others that occur stochastically, e.g. the remaining AA1 subclusters and the AA2 cluster. The validity of this hypothesis is strengthened by the high similarity of paired technical replicates, the lack of variance of the Ctrl expression phenotype over all 4 weeks, the existence of an AA1 subcluster populated by all AA replicates, and the shared and distinct stem cell- and CSC marker genes that were upregulated in AA1 and/or AA2.

### Acid-adapted pancreatic cancer cells exhibit aggressive growth and metastasis in vivo

To determine whether the increased stem cell potential of the acid-adapted pancreatic cancer cells would translate into increased aggressiveness *in vivo*, we orthotopically transplanted Ctrl and AA Panc-1 cells into the tail of the pancreas of NOD/SCID mice. In an initial experiment, we monitored body weight as a surrogate for tumor growth until a humane endpoint was reached (body weight below 80% of initial weight or signs of distress; Fig 6A, ‘humane endpoint’), after which we extracted pancreas, liver and lymph nodes (Fig. 6A). Body weight declined much more rapidly in the group injected with AA Panc-1 cells compared to the group injected with Ctrl Panc-1 cells (Fig. 6B, left panel), resulting in a significantly shorter survival (P=0.0159, Log-rank (Mantel-Cox) test) in the AA group (Fig. 6C). By visual inspection of the pancreata isolated at the humane endpoint it was evident that all mice presented with large tumors in the pancreas that were invading adjacent tissues. Moreover, macroscopic liver metastasis was apparent in the liver of both the Ctrl and the AA group.

**Figure 6.**
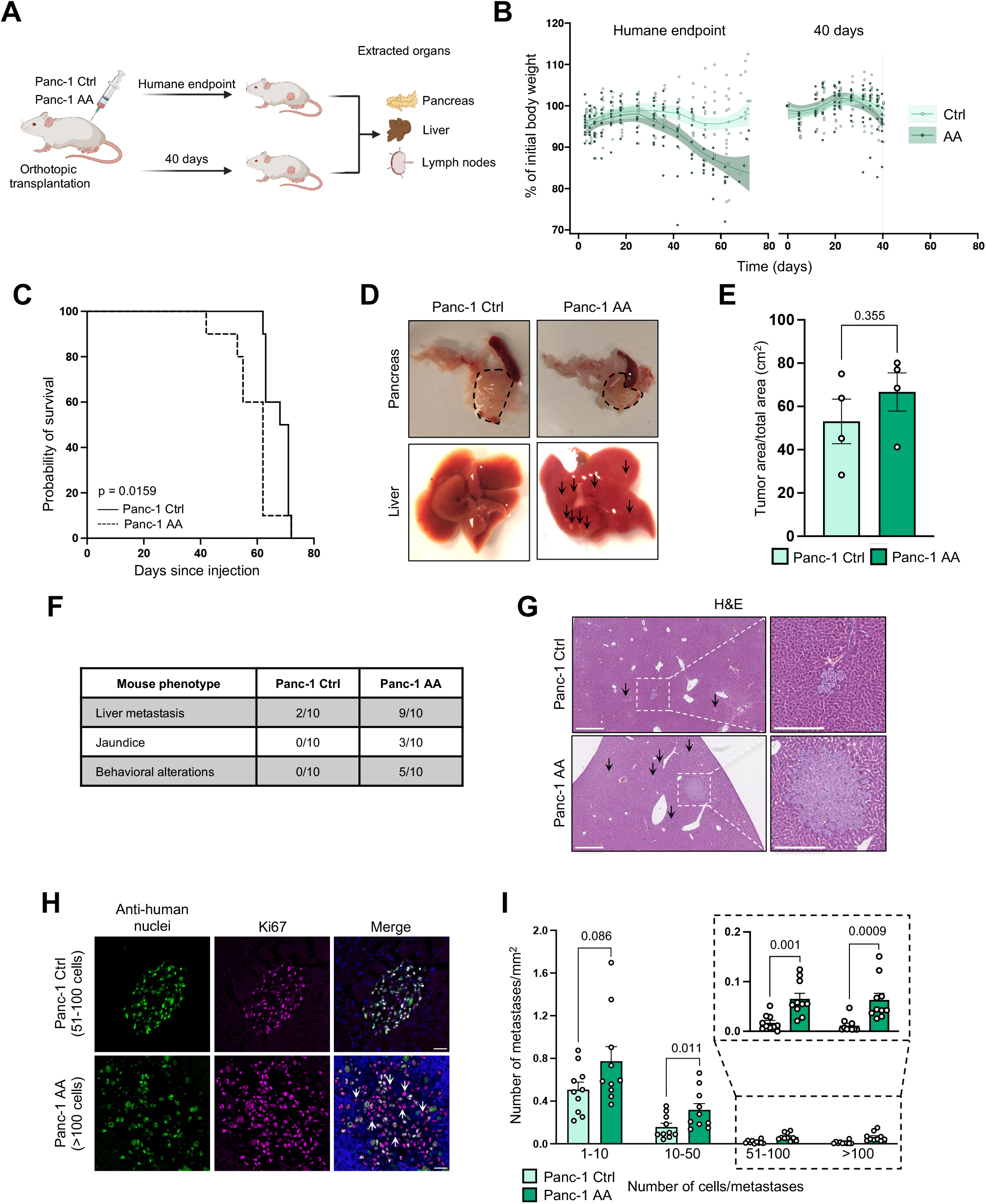
Acid adaptation accelerates PDAC progression in an orthotopic model. (**A**) Illustration of the *in vivo* experimental workflow. Two separate *in vivo* experiments were performed (‘humane endpoint’ and ‘40 days’, upper and lower arrows, respectively). In both experiments, Ctrl and AA Panc-1 cells were orthotopically transplanted into the tail of the pancreas of NOD/SCID mice and the pancreas, liver, and lymph nodes were extracted at the designated endpoint. In the humane endpoint experiment (upper arrow, panel (**B**-**C**)), the endpoint was determined as ≧20% weight loss or signs of severe illness, while in the time-based (lower arrow, panel (**B**, **D**-**I**)), the endpoint was 40 days. (**B**) Body weight percentage of initial for each mouse in the Ctrl and AA group (n=10 per group, per experiment) for the humane endpoint-(left) and time-based (right) *in vivo* experiments. Each dot represents one mouse, lines are a LOESS regression model with 95% confidence intervals and a smoothed function indicating the trend. (**C**) Kaplan-Meier survival curve of mice transplanted with Ctrl or AA Panc-1 cells. (**D**) Representative images of the pancreas and liver. Pancreatic tumor area is indicated by dotted lines and arrows denote macroscopic liver metastases. (**E**) Tumor area (cm^2^) determined from histopathological analysis as the fraction of tumor in the total pancreatic area. (**F**) Table reporting the number of mice with the indicated phenotype in each group at the endpoint in the time-based experiment. Mouse phenotypes include macroscopic liver metastases, jaundice and altered behavior (e.g. no self-grooming, no response to external stimuli). (**G**) Representative H&E-stained images of the liver showing metastasis in Ctrl and AA groups. Arrows denote metastases. Scale bar: 500 and 250 µm for full and zoom images, respectively. (**H**) Representative IF images of mice liver sections stained for anti-human nuclei (green), Ki67 (magenta) and nuclei (DAPI, blue). The upper and lower panel exemplifies metastases with 51-100 and >100 cells; respectively. Color intensity was adjusted independently for each image. White arrows in the lower merge image denote anti-human nuclei-negative, Ki67-positive cells. Scale bar: 20 µm. (**I**) Quantification of number of liver metastases per mm^2^ liver tissue divided into size groups (number of cells/metastases). For panel (**C**), a Log-rank (Mantel-Cox) test was performed, while in panels (**E**, **I**) a paired two-tailed student’s t-test was performed, AA vs Ctrl. In panel (**I**), the test was performed for each size group.

To allow direct comparison of tumor area and metastasis formation, we carried out a second experiment in which we orthotopically transplanted Panc-1 Ctrl and AA cells into NOD/SCID mice as above, but all mice were terminated at day 40 before any clinical manifestation of tumor growth, (see Fig. 6A, ‘40 days’). Because of the shorter growth period, there were no obvious differences in body weight, consistent with the pattern in this time period in the first experiment (Fig. 6B). Visual inspection of the pancreas and liver revealed that all animals presented with tumor growth in the pancreas (Fig. 6D, top), and liver metastasis appeared in both conditions, being more present in the AA group (Fig. 6D, bottom). Histopathological analysis of tumor area, determined as the fraction of tumor in the total pancreatic area (Fig. 6E), showed a slight, non-significant tendency for increased tumor area in the mice injected with AA cells compared to Ctrl cells. However, a striking difference was evident for the development of macroscopically visible metastases. In the AA group, 9 of the 10 mice presented with liver metastasis, 3 with jaundice at endpoint and 5 with behavioral alterations, compared to only two with liver metastasis in the control group (Fig. 6F).

Next, we assessed the extent of liver metastasis in Ctrl and AA cell-injected mice. Histopathological analysis revealed distinct masses growing in the livers of both groups (Fig. 6G). We employed immunofluorescence analysis using an antibody against the human-specific histone H3.3 variant to identify the presence of Panc-1 cells within the metastases at a single-cell resolution, and Ki67 staining to study the proliferative index of both human and murine cells (Fig. 6H). We categorized the metastatic lesions into four size groups based on the number of H3.3-positive (H3.3+) cells per mm^2^ (Fig. 6H-I; Suppl. Fig. 6). While the proliferative index of the metastatic lesions was similar in the Ctrl and AA groups (Suppl. Fig. 6), mice injected with AA Panc-1 cells developed significantly more metastases in the three largest sub-categories (10-50, 51-100, and >100 cells/metastasis) compared to mice injected with Ctrl Panc-1 cells. (Fig. 6I). Interestingly, in both groups, we also noted an infiltration of Ki-67-positive, H3.3-negative cells - i.e. proliferating murine cells - in the metastatic lesions (Fig. 6H, white arrows, Suppl. Fig. 6, 2^nd^ row, white arrow), suggesting an activation of murine stromal cells in the metastatic niche. This highlights a notable interaction between the human cancer cells and the mouse liver environment (Fig. 6H).

Collectively, these results show that acid adaptation of pancreatic cancer cells accelerates pancreatic cancer progression and increases metastatic potential *in vivo*.

## Discussion

We and others have demonstrated that the acidity of the tumor microenvironment favors several cancer cell traits associated with aggressive disease (Swietach *et al*, 2023; Corbet & Feron, 2017). These include increased 3D growth and adhesion-independent colony formation (Czaplinska *et al*, 2023), invasiveness and metastasis (Czaplinska *et al*, 2023)(Moellering *et al*, 2008)(Corbet *et al*, 2020) and profoundly rewired metabolism (Michl *et al*, 2022; Lamonte *et al*, 2013)(Corbet *et al*, 2014; Rolver *et al*, 2023). The key finding of the present work is that acid adaptation of pancreatic cancer cells drives upregulation of CSC properties and aggressive *in vivo* growth and metastasis.

Phenotypically, acid-adapted Panc-1 cells exhibited pronounced upregulation of ALDH activity, β-catenin activity, and pancreatosphere-forming capacity. These properties are strikingly characteristic of CSCs (Batlle & Clevers, 2017)(Vermeulen *et al*, 2008)(Hermann & Sainz, 2018). Our finding that acid adaptation strongly exacerbated the *in vivo* aggressiveness and metastatic potential of Panc-1 cells is also consistent with the notion that growth in the acidic tumor microenvironment selects for a population with CSC properties.

We did not identify a common acid adaptation-induced CSC marker profile between the human pancreatic cancer cell lines studied. This is not surprising given that a wide spectrum of different CSC profiles are reported in pancreatic cancer cells (Ishiwata *et al*, 2018)(Jaiswal *et al*, 2012; Kallifatidis *et al*, 2009)(Nimmakayala *et al*, 2021)(Li *et al*, 2007)(Wang *et al*, 2019)(Hermann & Sainz, 2018). Some, but not all, previously reported pancreatic cancer CSC genes were upregulated, and the Yamanaka factors tended to be downregulated. Thus, despite their classical functional CSC characteristics (ALDH activity, β-catenin activity, pancreatosphere-forming capacity, and, as previously demonstrated (Rolver *et al*, 2023), a shift toward OXPHOS-based metabolism, the acid adaptation-induced CSCs do not as an overall group exhibit a typical CSC marker profile. Strikingly, however, a strong upregulation of genes encoding the well-described pancreatic CSC markers CD9, cMET and EPCAM was revealed in one of the clusters identified by single nuclei sequencing of Panc-1 cells. The single nuclei sequencing revealed that acid adaptation resulted in several populations of cells, where some populations were captured in all replicates while some appeared stochastically. However, despite their differences, the acid-adapted clusters all showed upregulation of genes and GO terms associated with stemness and CSCs. Additionally, the acid-adapted cell clusters showed upregulation of marker genes associated with cancer aggressiveness, including EMT, metastasis, and resistance to cell death, many of which were shared between clusters.

The impact of chronic acidosis on stemness has, to our knowledge, not previously been studied in pancreatic cancer. However, it is well accepted that CSC properties are dependent on dedicated niches (Batlle & Clevers, 2017) and in the context of glioma, acidosis has been proposed to be an important property of this niche (Hjelmeland *et al*, 2011)(Filatova *et al*, 2016). Furthermore, it was recently shown that the ability of colon cancer cells to form lumen-containing colonies in matrigel correlated with their resistance to acid challenges, suggesting a link between CSC properties and capacity for growth under acidic conditions (Michl *et al*, 2024).

Which signaling events drive the acid-adaptation-induced increase in CSC population? Several mechanisms seem possible based on this and our previous work. Wnt/β-catenin signaling has been assigned a key role in driving pancreatic cancer CSC properties (Wu *et al*, 2019), and our findings suggest that acid-adapted pancreatic cancer cells exhibit upregulated Wnt signaling both via canonical (increased β-catenin activity) and noncanonical (p46JNK) pathways (Fig. 2; Suppl. Fig. 2B-C). Also ERK activity was increased in the acid-adapted cells. It is also interesting that the Yamanaka factor KLF4, which was downregulated in acid-adapted Panc-1 cells, is a negative regulator of pancreatic cancer stemness through regulation of CD44 (Yan *et al*, 2016).

Another interesting possibility is that the CSC properties of the acid-adapted cells are linked to the reported acid-induced shift toward increased reliance on oxidative phosphorylation (OXPHOS) and fatty acid β-oxidation (Rolver *et al*, 2023; Michl *et al*, 2022).

Increased dependence on OXPHOS is a general characteristic of CSCs including pancreatic CSCs (Sancho *et al*, 2015)(Courtois *et al*, 2021), and increased dependence on fatty acid β-oxidation has been demonstrated in CSCs from both breast (Wang *et al*, 2018) and pancreatic (Nimmakayala *et al*, 2021) cancer. In particular, upregulation of carnitine palmitoyltransferase 1B (CPT1B), which we recently showed to be driven by acid adaptation (Rolver *et al*, 2023), was linked to increased stemness in breast cancer (Wang *et al*, 2018). EEMT, shown to be increased by acid adaptation in several cell types (Corbet *et al*, 2020), has been assigned a role in pancreatic cancer CSCs (Batlle & Clevers, 2017)(Rodriguez-Aznar *et al*, 2019). Finally, the stress of acidic growth could select for a senescence-associated reprogramming which has also been associated with stemness (Milanovic *et al*, 2018).

Importantly, orthotopic transplantation experiments showed that, compared to Ctrl pancreatic cancer cells, acid-adapted cancer cells exhibited enhanced pancreatic tumor growth and metastasis *in vivo*, with a particularly dramatic difference in metastatic growth. Interestingly, no gross histopathological or morphological changes were observed in the primary tumors of acid adapted compared to control cells, suggesting that acid adaptation triggers cell-autonomous alterations in Panc-1 cells, enhancing their metastatic capability. Also the increased OXPHOS metabolism of acid-adapted cells could be relevant in this regard, as OXPHOS is linked to metastatic potential and may represent a relapse mechanism upon KRAS inhibition (Viale *et al*, 2014). Intriguingly, accumulating evidence suggests that OXPHOS metabolism might be a vulnerability in cancer cells, potentially serving as a basis to stratify patients for therapy or overcoming treatment resistance (Evans *et al*, 2021)(Ashton *et al*, 2018). This is in line with a recent CRISPR-Cas9 screen in colorectal cancer cells (Michl *et al*, 2022), which indicated that targeting OXPHOS could represent a novel therapeutic approach for cancers with high metastatic potential, particularly in acidic environments.

In conclusion, we report that adaptation of pancreatic cancer cells to growth at an extracellular pH in the range found in poorly perfused tumor regions selects for CSC properties, including increased ALDH- and β-catenin activity, pancreatosphere-forming efficiency, and increased metastatic growth *in vivo*. Although classical pancreatic cancer CSC populations were not overall increased by AA, single nucleus sequencing confirmed an acid adaptation-induced emergence of a stem cell population. We suggest that acid adaptation of pancreatic cancer cells enriches for a CSC population with enhanced self-renewal capacity, which could contribute importantly to the aggressiveness of pancreatic cancers. Our work provides new insight into the mechanisms through which the acidic tumor microenvironment favors aggressive cancer growth.

## Materials and Methods

### Cell lines, cell culture and acid adaptation

Panc-1, Patu8988S and MiaPaCa-2 human pancreatic cancer cells were cultured in RPMI (Sigma, #R1383) or DMEM (Gibco, #41966-029) adjusted to correct pH at 5% CO_2_ (7.6 and 6.5 for Ctrl and acid-adapted, respectively) either by titration with hydrochloric acid or by replacing the appropriate amount of NaHCO_3_^-^ with NaCl. Cell culture media were supplemented with 5-10% Fetal Bovine Serum (FBS, Gibco, #F9665), 1% Penicillin/Streptomycin (Sigma-Aldrich, #P0781) and for RPMI also 10 mM glucose (Sigma, #G8644). To ensure correct growth conditions, medium pH was measured inside the incubator, after equilibration at 5% CO_2_, using a mini-pH electrode. Acid-adaptation was carried out by culturing cells for 1 month in the pH 6.5 medium. Cells were cultured at 37°C, 95% humidity, 5% CO_2_, passaged at 70-80% confluency and discarded after passage 25.

### Gene set enrichment analysis

The Gene Set Enrichment Analysis (GSEA) pre-ranked function was used based on the log_2_fold change ranked acid adaptation gene list using the Broad Institute GSEA java tool version 4.0.3 or gene lists from (Wong *et al*, 2008). The gene sets GAL_LEUKEMIC_STEM_CELL_UP(M17428), BEIER_GLIOMA_STEM_CELL_UP (M9126), YAMASHITA_LIVER_CANCER_STEM_CELL_UP (M16956), MOUSE_ADULT_TISSUE_STEM_MODULE; CORE_ESC_LIKE_MODULE andHUMAN_ESC_LIKE_MODULE were used, all from (Wong *et al*, 2008). The number of permutations was set to 1000.

### ALDH activity

The ALDEFLUOR^TM^ Assay kit (Stem Cell Technologies, #01700) was used to determine ALDH activity according to manufacturers’ instructions. Briefly, cells were trypsinized and collected (500.000 cells/mL), followed by incubation with ALDEFLUOR^TM^ reagent (37°C, 45 min). Internal negative control samples were treated with a specific inhibitor of ALDH, diethylaminobenzaldehyde (DEAB), prior to incubation with ALDEFLUOR^TM^ reagent. ALDH activity was examined by flow cytometry using a Beckman Coulter CytoFLEX cytometer, followed by data analysis using FlowLogic software (Inivai Technologies). DEAB-treated samples were used to set the threshold for background fluorescence and define the ALDEFLUOR^TM^ positive gate (ALDH^+^) in DEAB-negative samples.

### SDS page and Western blotting

Western blot analyses were performed essentially as in (Rolver *et al*, 2023). Cells were grown to ∼80% confluency, washed in PBS and lysed (1% SDS, 10 mM Tris-HCl, 1 mM NaVO_3_, and protease inhibitors (Roche, #11836153001), pH 7.5, 95°C). Lysates were sonicated, centrifuged, and protein concentrations determined by Bio-Rad DC Protein assay (Bio-Rad, #500-0113, -0114, -0115). After equalizing protein concentrations with ddH_2_O, lysates were mixed with 1:1 NuPAGE LDS 4x Sample Buffer (Invitrogen, #NP0007) and 0.5 M dithiothreitol (Sigma #646563) and separated using 10% Criterion TGX Precast Midi Protein Gels (BioRad, #567-1035), Tris-Glycine-SDS running buffer (BioRad, #161-0732), and BenchMark ladder (Invitrogen, #10747-012). Proteins were transferred to nitrocellulose membranes (BioRad, #170-4159), stained with Ponceau S (Sigma-Aldrich, #7170-1L) and blocked (1 h, 37°C, 5% dry milk in TBST or 1% BSA (Sigma, #A7906)). Membranes were washed in TBST, incubated with primary antibodies (4°C, overnight (ON)), washed in TBST, incubated with HRP conjugated secondary antibodies (1 h, room temperature (RT)), washed in TBST, and developed using Clarity Western ECL Substrate (BioRad, #1705061). Band intensities were quantified using ImageJ and normalized to loading control. Primary and secondary antibodies are provided in Suppl. table 2.

### β-catenin reporter assay

Cells were seeded in 12-well plates and transfected at 50-70% confluence with 0.5 µg/well TOPflash (Addgene, #12456) and 0.3 µg pRL-TK (Promega) employing Lipofectamine 3000 (Invitrogen, #L3000015). After 4-5 h, medium was exchanged and after 48 h, lysates were acquired with Passive Lysis Buffer (Promega, #E1910), diluted in ddH_2_O, and luciferase activity assessed by Dual Luciferase Reporter Assay (Promega, #E1910) in a Varioskan LUX microplate reader (Thermo Fischer Scientific). Results are shown as fold increase in Firefly luciferase signal after background subtraction and normalization to pRL-TK Renilla luciferase signal.

### Pancreatosphere formation assay and quantification

#### First generation

Cells were grown to ∼70% confluency, washed, trypsinized, and resuspended in FBS-containing medium. Cell suspensions were centrifuged, the supernatant discarded, and the cells resuspended in 3-5 mL of pancreatosphere medium (DMEM/F12 (Gibco, #11320033), 1% P/S, EGF (20 ng/mL, Sigma, #E9644), FGF (20 ng/mL, R&D systems, #3718-FB-010), B27 (2%, Thermo Scientific, #12587010). Cells were dissociated to single cells by extensive pipetting or by using a 25 G needle, then counted, plated at 800 and/or 1600 cells/cm^2^ in ultra-low attachment wells (Corning, #3473) in 500 µL pancreatosphere medium, and incubated undisturbed for 5-7 days at 37°C and 5% CO_2_.

#### Second and third generation

For serial passaging, pancreatospheres were collected in tubes (independent of seeding density), centrifuged and incubated in 200 µL 0.5% trypsin (Sigma, #T4174) for 2-3 min at 37°C. Extensive pipetting or a 25 G needle was employed to create a single cell suspension. Hereafter, 400 µL FBS-containing medium was added, the tubes centrifuged and the cells resuspended in 200 µL pancreatosphere medium. Cells were counted, reseeded and incubated as above.

#### Data acquisition

Brightfield (BF) and fluorescence images of pancreatospheres (2-7 wells/condition) were acquired employing either an Olympus IX83 microscope with a Yokogawa scanning unit, 4X/0.13 PhL or 10X/0.3 Ph1 air objective, or a Nikon Ti2-E microscope with a Kinetix Teledyne Photometrics camera, Plan Fluor 4X/0.13 air objective. Nuclei were visualized by incubating pancreatospheres with hoechst (1:1000, 15 min, 37°C, Hoechst33342 (Thermo Fisher, #62249)) and acquiring images as z-stacks.

#### Data analysis

Due to differences in morphology between cell lines, the pancreatosphere-formation capacity was assessed either based on total number (PaTu) or size of pancreatospheres (Panc-1 and MiaPaCa-2). The area of Panc-1 pancreatospheres was determined by creating an imageJ macro (inspired from (Choudhry, 2016)). The macro detects, counts and measures pancreatospheres in a series of images automatically. Parameters such as minimum and maximum particle size were fixed across all images, whereas background subtraction was adjusted individually for accurate detection of pancreatosphere size. For MiaPaca-2 pancreatospheres, the freehand tool in ImageJ was used to determine the area of each pancreatosphere, while for Patu pancreatospheres, total number of pancreatospheres was counted and the pancreatosphere-forming efficiency (PFE) calculated as the number of pancreatospheres/well divided by the number of cells seeded/well. For each biological replicate, the average PFE was calculated and reported. For PaTu and MiaPaCa-2, the relative difference in PFE between Ctrl and AA was independent of seeding density and the acquired data has therefore been pooled. Z-projection and adjustments of brightness and contrast were performed as needed using ImageJ software.

### Flow cytometric analysis of stem cell populations

Cells were cultured to 70-80% confluency, trypsinized and collected (2x10^6^ cells per sample). Experimental and control samples (unstained/autofluorescence Ctrl, FMO Ctrl and compensation Ctrls) were centrifuged (3 min, 50 RCF) and resuspended in flow buffer (PBS, 2% FBS, and 0.1% NaN_3_). Cells were labeled by adding 100 µL of flow buffer mixed with relevant antibodies and incubated for 30 min at 4°C. Subsequently, cells were fixed (30 min, 4°C, PBS containing 2 % FBS and 1% PFA), resuspended in flow buffer, transferred to flow cytometry tubes (5 mL Round-Bottom Polystyrene tubes, Falcon, #352058), and kept at 4°C until analysis. Samples were analyzed using either a BD FACS Calibur or a Beckton Coulter Cytoflex. Data processing was performed using FlowLogic v. 8.6 (Inivai Technologies). Antibodies are provided in Supplementary table 2.

### Single nucleus preparation, sequencing and analysis

#### Nuclei isolation

Nuclei were isolated from Panc-1 cells using an IgePal-based lysis buffer (protocol optimized from (Krishnaswami *et al*, 2016)), see Supplemental Material and methods for further details). Briefly, 4x10^6^ cells were collected in BSA-coated DNA LoBind tubes (Eppendorf, #022431021), lysed over ice (10 min) and centrifuged (350 RCF, 5 min, 4°C). Lysed cells/nuclei were washed in ice-cold PBS supplemented with BSA (3%), Protease and RNAse inhibitors, followed by centrifugation (800 RCF, 10 min, 4°C). This wash step was repeated twice. For optimal removal of cell membrane, nuclei were spun down through a gradient of Iodixanol (50 % over 29%, 14000 RCF, 22 min, “Optiprep”, Stemcell Cat #7820), washed and filtered through 30 µm filters (Miltenyi Biotec, #130-098-458). Nuclei isolation was verified by BF imaging, and a control sample was stained with 4’,6-diamidino-2-phenylindole (DAPI) to assess nuclear membrane integrity. Nuclei were counted and adjusted to a final concentration of 700-1200 nuclei/µL in nuclei storage buffer.

#### Generation of cDNA libraries and sequencing

cDNA libraries were prepared using 10X Genomics Chromium technology at the Single Cell Genomics Core Facility at the Biotech Research & Innovation Centre, University of Copenhagen. Briefly, 5-10*10^4^ nuclei along with reagents, and partitioning oil were loaded onto the 10X Chromium chip (3’ V3.1) according to manufacturer’s recommendations (Chromium Next GEM Single Cell 3ʹ Reagent Kits v3.1, Document Number CG000204 Rev D, 10x Genomics), and processed in the 10X Chromium controller (10X Genomics). This generated Gel Beads in Emulsion (GEMs) droplets containing single nuclei, a barcoded gel bead and reagents for reverse transcription. Inside the GEMs, nuclei were lysed, and the mRNA reverse transcribed into full-length cDNA with a barcoded cell code and a unique molecular identifier (UMI). Hereafter, cDNA was released, cleaned up, amplified, fragmented, and a sample index attached according to manufacturer’s recommendations. Resulting single nuclei cDNA libraries were pooled and diluted to 1.5 nM, followed by sequencing at the Core Facility for Flow Cytometry and Single Cell Analysis, Faculty of Health and Medical Sciences, University of Copenhagen on a NovaSeq 6000 (Illumina, Inc.) using the NovaSeq S2 sequencing kit v. 1.5 (100 cycles)(Illumina, Inc., #20028316) according to manufacturer’s recommendations (NovaSeq 6000 Denature and Dilute Libraries Guide, document no. 1000000106351 v03, and NovaSeq 6000 System Guide, document no. 1000000019358 v17, Illumina, Inc.).

#### Computational analysis

Raw reads in BCL files were demultiplexed, FASTQ-converted and mapped to human transcriptome GRCh38 using CellRanger (7.2.0: https://www.10xgenomics.com/ support/software/cell-ranger/latest). Ambient RNA was removed through CellBender (0.3.1) (Fleming *et al*, 2023). The resulting count matrices were quality filtered in Seurat (5.0.1) (Hao *et al*, 2024). Cells with less than 1000 genes detected or with mitochondrial RNA% > 10% were filtered out. Non-protein-coding genes were removed before normalization and scaling. UMAPs were constructed based on the PCA of the scaled data (number of components=30). Across all 8 samples, we identified 30 cell clusters via Louvain clustering (res=0.8). From those clusters, three major cell populations were discovered. Differentially expressed genes across cell populations were identified using FindMarkers function in Seurat (Wilcoxon tests, FDR adjusted). Resulting differentially expressed genes were subjected to gene ontology enrichment analysis via clusterProfiler (Wu *et al*, 2021) (4.6.2, all expressed genes as enrichment background). Cell cycle phase of each cell was predicted using the cyclone function from scran package (Lun *et al*, 2016) (1.26.2). Wilcoxon rank sum test was used to assess the change of proportion of G0/G1 phase cells across cell groups. R packages ggplot2 (Wickham, 2009) (3.5.1) and SingleCellPipeline (0.5.1;https://github.com/zhanghao-njmu/SCP) were applied to visualize the results.

### Orthotopic tumor model

Orthotopic transplantations were performed using NOD/SCID mice (MGI:2163032). Buprenorphine was administered subcutaneously (0.05-0.1 mg/kg) prior to the initial incision in the back of the animal and another dose up to 2 h after surgery if signs of pain were still visible. Mice were anesthetized by isoflurane inhalation (2-2.5%). The surgical site was shaved and disinfected with 70% ethanol. A horizontal laparotomy of approx. 1 cm at the location of the spleen was performed through the skin, followed by the muscle layer. The pancreas was made visible by pulling gently on the spleen. Cells were prepared at a concentration of 20x10^6^ cells/ml Matrigel®, and 50 µl of Matrigel (10^6^ cells) were injected in the tail/body of the pancreas using BD Ultrafine™ 8mmX30G syringes. The needle was left in place for 5 s to avoid reflux of the injected volume. The pancreas and spleen were gently placed back again in the abdominal cavity. Wound closure of the muscle layer and skin was performed separately through the continuous suture technique. Absorbable sutures (4-0 or 3-0 nylon or vicryl) were used for wound closure. Animals were housed in accordance with best animal husbandry guideline recommendations of the European Union Directive (2010/63/EU). The Danish Animal Experiments Inspectorate reviewed and approved all animal experiments. The animal license number for these experiments is 2019-15-0202-00183.

### Organ extraction

Once the animal was euthanized, a vertical incision from the neck to the lower part of the abdomen was performed to expose the ventral cavity. Two incisions were made in the skin along both legs to expose the inguinal lymph nodes attached to the internal side. The liver was isolated in the ventral cavity and detached from the stomach before removing the connective tissue around it. The pancreas first detached from the colon until reaching the duodenum. To continue, we horizontally sectioned the tissue attached along the duodenum from left to right until the lower part of the stomach. Finally, we removed all connective tissue attaching it to the spleen and the ventral cavity.

### H&E and Immunofluorescence stainings

Pancreatic tissues were fixed using 4% paraformaldehyde (VWR Chemicals, #9713-1000) for 24 h, then dehydrated in 70% ethanol (VWR Chemicals, #20824.365) at RT, and embedded in paraffin. The tissues were cut into 4-10 μm sections and placed on Superfrost Plus slide (10149870; Fisher Scientific). Hematoxylin and eosin (H&E) staining was performed using the H&E Staining Kit (Abcam, Cambridge, UK) following the manufacturer’s instructions. Whole-section imaging was conducted using a NanoZoomer-XR Digital Slide Scanner C12000-01 (Hamamatsu).

For immunofluorescence analysis, antigen retrieval was performed with Tris-EDTA buffer (pH 9). Hereafter, the sections were washed in PBS-Tween 0.5%, blocked with 1% donkey serum, and incubated with primary antibodies (4°C, overnight). The following day, the sections were incubated with secondary antibodies (1 h, room temperature) and mounted with Vectashield® Mounting Medium (Vector Laboratories, #H-1000). Image acquisition was performed using a Leica SP8 confocal microscope, with consistent laser intensities and parameters, a HC Plan-Apochromat 40x / 1.1 W objective and the following detectors: 2 x super-sensitive Hybrid detectors (HyD) and 1 x T-PMT detector. A list of antibodies is provided in Supplementary table 2.

## Data analysis and statistics

Data is shown as representative images, or as mean measurements with standard error of the mean (SEM) error bars. Points in bar and box plots show biological experiments. Technical replicates are given in figure legends where relevant. The statistical analyses were performed using GraphPad Prism version 10.0 and R version 4.2.2. To test for statistically significant differences between two groups, a two-tailed Student’s *t*-test was performed. In the Kaplan-Meier survival curve, the P-value was calculated using the Log-rank (Mantel-Cox) test. P-value threshold for significance was set as 0.05, and exact p-values are shown in figures. Additional data analysis and illustrations were performed using Microsoft Excel, GraphPad Prism, ImageJ and BioRender.com. Single nuclei RNA-seq analysis was made using Seurat and clusterProfiler, including statistical tests (see relevant methods section above).

## Author Contributions

Conceptualization, MR, SFP, LA, AS; methodology, RD, NSP, MF, JB, YD, EA; formal analysis, RD, NSP, JY, RI, MR, MF, JCR, YD, AS, SFP; investigation, MR, JCR, YD, MF, RI, JH, RD, NSP, AB, JB, JY, AS; data curation, MR, MF, YD; writing—original draft preparation, SFP; writing—review and editing, MR, JCR, YD, AS, LA, SFP; visualization, MR, JCR, JY, YD, AS; supervision, MR, MF, JB, AS, LA, SFP; project administration, MR, AS, LA, SFP; funding acquisition, MR, AS, LA, SFP. All authors have read and agreed to the published version of the manuscript.

## Data Availability Statement

The snRNA-seq data is available at the GEO database under accession number GSE269521 (for referees: please use the token qlebaeugbdwzjyp). Other data, including computer code for R and ImageJ analyses, are available from the corresponding author upon reasonable request.

## Acknowledgments

We gratefully acknowledge the excellent technical assistance of Tanja Larsen and Frida J. Birkbak, and contributions to early experiments by Eliana Alves, Muthulakshmi Ponniah, Josephine S. Kapel, Lukas K. Laursen, Anne B. Olsen and Naiyee R. Toiviainen. We are grateful to Irina Korshunova and Konstantin Khodosevich at the BRIC single cell genomics facility for excellent guidance and technical assistance.

## References

Ashton TM, McKenna WG, Kunz-Schughart LA & Higgins GS (2018) Oxidative Phosphorylation as an Emerging Target in Cancer Therapy. Clin Cancer Res 24: 2482– 2490

Batlle E & Clevers H (2017) Cancer stem cells revisited. Nat Med 23: 1124–1134

Boedtkjer E & Pedersen SF (2020) The Acidic Tumor Microenvironment as a Driver of Cancer. Annu Rev Physiol 82: 103–126

Choudhry P (2016) High-Throughput Method for Automated Colony and Cell Counting by Digital Image Analysis Based on Edge Detection. PLoS One 11: e0148469

Corbet C, Bastien E, Santiago de Jesus JP, Dierge E, Martherus R, Vander Linden C, Doix B, Degavre C, Guilbaud C, Petit L, et al (2020) TGFβ2-induced formation of lipid droplets supports acidosis-driven EMT and the metastatic spreading of cancer cells. Nat Commun 11: 454

Corbet C, Draoui N, Polet F, Pinto A, Drozak X, Riant O & Feron O (2014) The SIRT1/HIF2α axis drives reductive glutamine metabolism under chronic acidosis and alters tumor response to therapy. Cancer Res 74: 5507–5519

Corbet C & Feron O (2017) Tumour acidosis: from the passenger to the driver’s seat. Nat Rev Cancer 17: 577–593

Courtois S, de Luxán-Delgado B, Penin-Peyta L, Royo-García A, Parejo-Alonso B, Jagust P, Alcalá S, Rubiolo JA, Sánchez L, Sainz B Jr, et al (2021) Inhibition of Mitochondrial Dynamics Preferentially Targets Pancreatic Cancer Cells with Enhanced Tumorigenic and Invasive Potential. Cancers 13

Czaplinska D, Ialchina R, Andersen HB, Yao J, Stigliani A, Dannesboe J, Flinck M, Chen X, Mitrega J, Gnosa SP, et al (2023) Crosstalk between tumor acidosis, p53 and extracellular matrix regulates pancreatic cancer aggressiveness. Int J Cancer 152: 1210–1225

Davis JE Jr, Kirk J, Ji Y & Tang DG (2019) Tumor Dormancy and Slow-Cycling Cancer Cells. Adv Exp Med Biol 1164: 199–206

Dierge E, Debock E, Guilbaud C, Corbet C, Mignolet E, Mignard L, Bastien E, Dessy C, Larondelle Y & Feron O (2021) Peroxidation of n-3 and n-6 polyunsaturated fatty acids in the acidic tumor environment leads to ferroptosis-mediated anticancer effects. Cell Metab 33: 1701–1715.e5

Eser S, Reiff N, Messer M, Seidler B, Gottschalk K, Dobler M, Hieber M, Arbeiter A, Klein S, Kong B, et al (2013) Selective requirement of PI3K/PDK1 signaling for Kras oncogene-driven pancreatic cell plasticity and cancer. Cancer Cell 23: 406–420

Evans KW, Yuca E, Scott SS, Zhao M, Paez Arango N, Cruz Pico CX, Saridogan T, Shariati M, Class CA, Bristow CA, et al (2021) Oxidative Phosphorylation Is a Metabolic Vulnerability in Chemotherapy-Resistant Triple-Negative Breast Cancer. Cancer Res 81: 5572–5581

Filatova A, Seidel S, Böğürcü N, Gräf S, Garvalov BK & Acker T (2016) Acidosis Acts through HSP90 in a PHD/VHL-Independent Manner to Promote HIF Function and Stem Cell Maintenance in Glioma. Cancer Res 76: 5845–5856

Fleming SJ, Chaffin MD, Arduini A, Akkad A-D, Banks E, Marioni JC, Philippakis AA, Ellinor PT & Babadi M (2023) Unsupervised removal of systematic background noise from droplet-based single-cell experiments using CellBender. Nat Methods 20: 1323–1335

GBD 2017 Pancreatic Cancer Collaborators (2019) The global, regional, and national burden of pancreatic cancer and its attributable risk factors in 195 countries and territories, 1990-2017: a systematic analysis for the Global Burden of Disease Study 2017. Lancet Gastroenterol Hepatol 4: 934–947

Gires O, Pan M, Schinke H, Canis M & Baeuerle PA (2020) Expression and function of epithelial cell adhesion molecule EpCAM: where are we after 40 years? Cancer Metastasis Rev 39: 969–987

Hao Y, Stuart T, Kowalski MH, Choudhary S, Hoffman P, Hartman A, Srivastava A, Molla G, Madad S, Fernandez-Granda C, et al (2024) Dictionary learning for integrative, multimodal and scalable single-cell analysis. Nat Biotechnol 42: 293–304

Hermann PC, Huber SL, Herrler T, Aicher A, Ellwart JW, Guba M, Bruns CJ & Heeschen C (2007) Distinct populations of cancer stem cells determine tumor growth and metastatic activity in human pancreatic cancer. Cell Stem Cell 1: 313–323

Hermann PC & Sainz B Jr (2018) Pancreatic cancer stem cells: A state or an entity? Semin Cancer Biol 53: 223–231

Higgins DMO, Caliva M, Schroeder M, Carlson B, Upadhyayula PS, Milligan BD, Cheshier SH, Weissman IL, Sarkaria JN, Meyer FB, et al (2020) Semaphorin 3A mediated brain tumor stem cell proliferation and invasion in EGFRviii mutant gliomas. BMC Cancer 20: 1213

Hjelmeland AB, Wu Q, Heddleston JM, Choudhary GS, MacSwords J, Lathia JD, McLendon R, Lindner D, Sloan A & Rich JN (2011) Acidic stress promotes a glioma stem cell phenotype. Cell Death Differ 18: 829–840

Ishiwata T, Matsuda Y, Yoshimura H, Sasaki N, Ishiwata S, Ishikawa N, Takubo K, Arai T & Aida J (2018) Pancreatic cancer stem cells: features and detection methods. Pathol Oncol Res 24: 797–805

Jaiswal KR, Xin H-W, Anderson A, Wiegand G, Kim B, Miller T, Hari D, Ray S, Koizumi T, Rudloff U, et al (2012) Comparative testing of various pancreatic cancer stem cells results in a novel class of pancreatic-cancer-initiating cells. Stem Cell Res 9: 249–260

Jang G, Oh J, Jun E, Lee J, Kwon JY, Kim J, Lee S-H, Kim SC, Cho S-Y & Lee C (2022) Direct cell-to-cell transfer in stressed tumor microenvironment aggravates tumorigenic or metastatic potential in pancreatic cancer. NPJ Genom Med 7: 63

Jiao Y, Li Y, Liu S, Chen Q & Liu Y (2019) serves as a diagnostic and prognostic biomarker for pancreatic cancer. Onco Targets Ther 12: 4141–4152

Kallifatidis G, Rausch V, Baumann B, Apel A, Beckermann BM, Groth A, Mattern J, Li Z, Kolb A, Moldenhauer G, et al (2009) Sulforaphane targets pancreatic tumour-initiating cells by NF-kappaB-induced antiapoptotic signalling. Gut 58: 949–963

Kimbrough CW, Khanal A, Zeiderman M, Khanal BR, Burton NC, McMasters KM, Vickers SM, Grizzle WE & McNally LR (2015) Targeting Acidity in Pancreatic Adenocarcinoma: Multispectral Optoacoustic Tomography Detects pH-Low Insertion Peptide Probes In Vivo. Clin Cancer Res 21: 4576–4585

Kim MP, Fleming JB, Wang H, Abbruzzese JL, Choi W, Kopetz S, McConkey DJ, Evans DB & Gallick GE (2011) ALDH activity selectively defines an enhanced tumor-initiating cell population relative to CD133 expression in human pancreatic adenocarcinoma. PLoS One 6: e20636

Krishnaswami SR, Grindberg RV, Novotny M, Venepally P, Lacar B, Bhutani K, Linker SB, Pham S, Erwin JA, Miller JA, et al (2016) Using single nuclei for RNA-seq to capture the transcriptome of postmortem neurons. Nat Protoc 11: 499–524

Lamonte G, Tang X, Chen JL-Y, Wu J, Ding C-KC, Keenan MM, Sangokoya C, Kung H-N, Ilkayeva O, Boros LG, et al (2013) Acidosis induces reprogramming of cellular metabolism to mitigate oxidative stress. Cancer Metab 1: 23

Lee CJ, Dosch J & Simeone DM (2008) Pancreatic cancer stem cells. J Clin Oncol 26: 2806–2812

Li C, Heidt DG, Dalerba P, Burant CF, Zhang L, Adsay V, Wicha M, Clarke MF & Simeone DM (2007) Identification of pancreatic cancer stem cells. Cancer Res 67: 1030–1037

Lien W-H & Fuchs E (2014) Wnt some lose some: transcriptional governance of stem cells by Wnt/β-catenin signaling. Genes Dev 28: 1517–1532

Liu M, Zhang Y, Yang J, Zhan H, Zhou Z, Jiang Y, Shi X, Fan X, Zhang J, Luo W, et al (2021) Zinc-Dependent Regulation of ZEB1 and YAP1 Coactivation Promotes Epithelial-Mesenchymal Transition Plasticity and Metastasis in Pancreatic Cancer. Gastroenterology 160: 1771–1783.e1

Liu Y, Reyes E, Castillo-Azofeifa D, Klein OD, Nystul T & Barber DL (2023) Intracellular pH dynamics regulates intestinal stem cell lineage specification. Nat Commun 14: 3745

Li W, Wang Z, Zha L, Kong D, Liao G & Li H (2017) HMGA2 regulates epithelial-mesenchymal transition and the acquisition of tumor stem cell properties through TWIST1 in gastric cancer. Oncol Rep 37: 185–192

Lun ATL, McCarthy DJ & Marioni JC (2016) A step-by-step workflow for low-level analysis of single-cell RNA-seq data with Bioconductor. F1000R*es* 5: 2122

Ma Z, Li Z, Wang S, Zhou Q, Ma Z, Liu C, Huang B, Zheng Z, Yang L, Zou Y, et al (2021) SLC39A10 Upregulation Predicts Poor Prognosis, Promotes Proliferation and Migration, and Correlates with Immune Infiltration in Hepatocellular Carcinoma. J Hepatocell Carcinoma 8: 899–912

Metcalfe C & Bienz M (2011) Inhibition of GSK3 by Wnt signalling--two contrasting models. J Cell Sci 124: 3537–3544

Michl J, Wang Y, Monterisi S, Blaszczak W, Beveridge R, Bridges EM, Koth J, Bodmer WF & Swietach P (2022) CRISPR-Cas9 screen identifies oxidative phosphorylation as essential for cancer cell survival at low extracellular pH. Cell Rep 38: 110493

Michl J, White B, Monterisi S, Bodmer WF & Swietach P (2024) Phenotypic screen of sixty-eight colorectal cancer cell lines identifies CEACAM6 and CEACAM5 as markers of acid resistance. Proc Natl Acad Sci U S A 121: e2319055121

Milanovic M, Fan DNY, Belenki D, Däbritz JHM, Zhao Z, Yu Y, Dörr JR, Dimitrova L, Lenze D, Monteiro Barbosa IA, et al (2018) Senescence-associated reprogramming promotes cancer stemness. Nature 553: 96–100

Miyai T, Hojyo S, Ikawa T, Kawamura M, Irié T, Ogura H, Hijikata A, Bin B-H, Yasuda T, Kitamura H, et al (2014) Zinc transporter SLC39A10/ZIP10 facilitates antiapoptotic signaling during early B-cell development. Proc Natl Acad Sci U S A 111: 11780–11785

Moellering RE, Black KC, Krishnamurty C, Baggett BK, Stafford P, Rain M, Gatenby RA & Gillies RJ (2008) Acid treatment of melanoma cells selects for invasive phenotypes. Clin Exp Metastasis 25: 411–425

Nallasamy P, Nimmakayala RK, Karmakar S, Leon F, Seshacharyulu P, Lakshmanan I, Rachagani S, Mallya K, Zhang C, Ly QP, et al (2021) Pancreatic Tumor Microenvironment Factor Promotes Cancer Stemness via SPP1-CD44 Axis. Gastroenterology 161: 1998–2013.e7

Neoptolemos JP, Kleeff J, Michl P, Costello E, Greenhalf W & Palmer DH (2018) Therapeutic developments in pancreatic cancer: current and future perspectives. Nat Rev Gastroenterol Hepatol 15: 333–348

Nimmakayala RK, Leon F, Rachagani S, Rauth S, Nallasamy P, Marimuthu S, Shailendra GK, Chhonker YS, Chugh S, Chirravuri R, et al (2021) Metabolic programming of distinct cancer stem cells promotes metastasis of pancreatic ductal adenocarcinoma. Oncogene 40: 215–231

Patil K, Khan FB, Akhtar S, Ahmad A & Uddin S (2021) The plasticity of pancreatic cancer stem cells: implications in therapeutic resistance. Cancer Metastasis Rev 40: 691–720

Pedersen SF & Counillon L (2019) The SLC9A-C Mammalian Na/H Exchanger Family: Molecules, Mechanisms, and Physiology. Physiol Rev 99: 2015–2113

Pedersen SF, Novak I, Alves F, Schwab A & Pardo LA (2017) Alternating pH landscapes shape epithelial cancer initiation and progression: Focus on pancreatic cancer. Bioessays 39

Plaks V, Kong N & Werb Z (2015) The cancer stem cell niche: how essential is the niche in regulating stemness of tumor cells? Cell Stem Cell 16: 225–238

Pothula SP, Xu Z, Goldstein D, Pirola RC, Wilson JS & Apte MV (2020) Targeting HGF/c-MET Axis in Pancreatic Cancer. Int J Mol Sci 21

Raggi C, Taddei ML, Sacco E, Navari N, Correnti M, Piombanti B, Pastore M, Campani C, Pranzini E, Iorio J, et al (2021) Mitochondrial oxidative metabolism contributes to a cancer stem cell phenotype in cholangiocarcinoma. J Hepatol 74: 1373–1385

Ren X, Feng C, Wang Y, Chen P, Wang S, Wang J, Cao H, Li Y, Ji M & Hou P (2023) SLC39A10 promotes malignant phenotypes of gastric cancer cells by activating the CK2-mediated MAPK/ERK and PI3K/AKT pathways. Exp Mol Med 55: 1757–1769

Rodriguez-Aznar E, Wiesmüller L, Sainz B Jr & Hermann PC (2019) EMT and stemness-key players in pancreatic cancer stem cells. Cancers 11: 1136

Rojas LA, Sethna Z, Soares KC, Olcese C, Pang N, Patterson E, Lihm J, Ceglia N, Guasp P, Chu A, et al (2023) Personalized RNA neoantigen vaccines stimulate T cells in pancreatic cancer. Nature 618: 144–150

Rolver MG, Holland LKK, Ponniah M, Prasad NS, Yao J, Schnipper J, Kramer S, Elingaard-Larsen L, Pedraz-Cuesta E, Liu B, et al (2023) Chronic acidosis rewires cancer cell metabolism through PPARα signaling. Int J Cancer 152: 1668–1684

Sancho P, Burgos-Ramos E, Tavera A, Bou Kheir T, Jagust P, Schoenhals M, Barneda D, Sellers K, Campos-Olivas R, Graña O, et al (2015) MYC/PGC-1α Balance Determines the Metabolic Phenotype and Plasticity of Pancreatic Cancer Stem Cells. Cell Metab 22: 590–605

Swietach P, Boedtkjer E & Pedersen SF (2023) How protons pave the way to aggressive cancers. Nat Rev Cancer 23: 825–841

Takahashi K & Yamanaka S (2006) Induction of pluripotent stem cells from mouse embryonic and adult fibroblast cultures by defined factors. Cell 126: 663–676

Vermeulen L, Sprick MR, Kemper K, Stassi G & Medema JP (2008) Cancer stem cells--old concepts, new insights. Cell Death Differ 15: 947–958

Viale A, Pettazzoni P, Lyssiotis CA, Ying H, Sánchez N, Marchesini M, Carugo A, Green T, Seth S, Giuliani V, et al (2014) Oncogene ablation-resistant pancreatic cancer cells depend on mitochondrial function. Nature 514: 628–632

Vlashi E, Lagadec C, Vergnes L, Matsutani T, Masui K, Poulou M, Popescu R, Della Donna L, Evers P, Dekmezian C, et al (2011) Metabolic state of glioma stem cells and nontumorigenic cells. Proc Natl Acad Sci U S A 108: 16062–16067

Wang L, Dong P, Wang W, Huang M & Tian B (2017) Gemcitabine treatment causes resistance and malignancy of pancreatic cancer stem-like cells via induction of lncRNA HOTAIR. Exp Ther Med 14: 4773–4780

Wang T, Fahrmann JF, Lee H, Li Y-J, Tripathi SC, Yue C, Zhang C, Lifshitz V, Song J, Yuan Y, et al (2018) JAK/STAT3-Regulated Fatty Acid β-Oxidation Is Critical for Breast Cancer Stem Cell Self-Renewal and Chemoresistance. Cell Metab 27: 136–150.e5

Wang VM-Y, Ferreira RMM, Almagro J, Evan T, Legrave N, Zaw Thin M, Frith D, Carvalho J, Barry DJ, Snijders AP, et al (2019) CD9 identifies pancreatic cancer stem cells and modulates glutamine metabolism to fuel tumour growth. Nat Cell Biol 21: 1425–1435

Wang YJ, Bailey JM, Rovira M & Leach SD (2013) Sphere-forming assays for assessment of benign and malignant pancreatic stem cells. Methods Mol Biol 980: 281–290

Wickham H (2009) ggplot2: elegant graphics for data analysis New York: Springer

Wong DJ, Liu H, Ridky TW, Cassarino D, Segal E & Chang HY (2008) Module map of stem cell genes guides creation of epithelial cancer stem cells. Cell Stem Cell 2: 333–344

Wu H-Y, Yang M-C, Ding L-Y, Chen CS & Chu P-C (2019) p21-Activated kinase 3 promotes cancer stem cell phenotypes through activating the Akt-GSK3β-β-catenin signaling pathway in pancreatic cancer cells. Cancer Lett 456: 13–22

Wu M, Zhang X, Zhang W, Chiou YS, Qian W, Liu X, Zhang M, Yan H, Li S, Li T, et al (2022) Cancer stem cell regulated phenotypic plasticity protects metastasized cancer cells from ferroptosis. Nat Commun 13: 1371

Wu T, Hu E, Xu S, Chen M, Guo P, Dai Z, Feng T, Zhou L, Tang W, Zhan L, et al (2021) clusterProfiler 4.0: A universal enrichment tool for interpreting omics data. Innovation (Camb) 2: 100141

Xu X, Chai S, Wang P, Zhang C, Yang Y, Yang Y & Wang K (2015) Aldehyde dehydrogenases and cancer stem cells. Cancer Lett 369: 50–57

Xu Z, Goel HL, Burkart C, Burman L, Chong YE, Barber AG, Geng Y, Zhai L, Wang M, Kumar A, et al (2023) Inhibition of VEGF binding to neuropilin-2 enhances chemosensitivity and inhibits metastasis in triple-negative breast cancer. Sci Transl Med 15: eadf1128

Yan Y, Li Z, Kong X, Jia Z, Zuo X, Gagea M, Huang S, Wei D & Xie K (2016) KLF4-Mediated Suppression of CD44 Signaling Negatively Impacts Pancreatic Cancer Stemness and Metastasis. Cancer Res 76: 2419–2431

Yan Y, Teng H, Hang Q, Kondiparthi L, Lei G, Horbath A, Liu X, Mao C, Wu S, Zhuang L, et al (2023) SLC7A11 expression level dictates differential responses to oxidative stress in cancer cells. Nat Commun 14: 3673

Yao J, Czaplinska D, Ialchina R, Schnipper J, Liu B, Sandelin A & Pedersen SF (2020) Cancer Cell Acid Adaptation Gene Expression Response Is Correlated to Tumor-Specific Tissue Expression Profiles and Patient Survival. Cancers 12

Zhao Y, Qin C, Zhao B, Wang Y, Li Z, Li T, Yang X & Wang W (2023) Pancreatic cancer stemness: dynamic status in malignant progression. J Exp Clin Cancer Res 42: 122

Zhou S, Schuetz JD, Bunting KD, Colapietro AM, Sampath J, Morris JJ, Lagutina I, Grosveld GC, Osawa M, Nakauchi H, et al (2001) The ABC transporter Bcrp1/ABCG2 is expressed in a wide variety of stem cells and is a molecular determinant of the side-population phenotype. Nat Med 7: 1028–1034

